# Two splice forms of *OsbZIP1*, a homolog of *AtHY5*, function to regulate skoto- and photo-morphogenesis in rice

**DOI:** 10.1101/2023.01.30.526072

**Authors:** Akanksha Bhatnagar, Naini Burman, Eshan Sharma, Akhilesh Tyagi, Paramjit Khurana, Jitendra P. Khurana

## Abstract

Plants possess well-developed light sensing mechanisms and signal transduction systems for regulating photomorphogenesis. ELONGATED HYOCOTYL 5 (HY5), a basic leucine zipper transcription factor, has been extensively characterized in dicot plants. In this study, we have shown that *OsbZIP1* is a functional homolog of *Arabidopsis HY5 (AtHY5)* and is important for light-mediated regulation of seedling and mature plant development in rice. Ectopic expression of *OsbZIP1* in rice reduces plant height and leaf length without affecting plant fertility, which is in contrast to *OsbZIP48*, another HY5 homolog we characterised earlier. *OsbZIP1* is alternatively spliced and the isoform OsbZIP1.2 lacking COP1 binding domain regulates seedling development in dark; this is unique since AtHY5 lacking COP1 binding domain does not display such a phenotype. Rice seedlings overexpressing *OsbZIP1* were found to be shorter than vector control under white and monochromatic light conditions whereas RNAi seedlings displayed completely opposite phenotype. While OsbZIP1.1 is light regulated, OsbZIP1.2 shows similar protein profile in both light and dark conditions. Due to its interaction with OsCOP1, OsbZIP1.1 undergoes degradation via 26S proteasome under dark conditions. Also, OsbZIP1.1 interacts with CASEIN KINASE 2 (OsCK2ɑ3) and consequently undergoes phosphorylation. In comparison, OsbZIP1.2 did not show any interaction with COP1 and OsCK2ɑ3. We propose that OsbZIP1.1 most likely works under low fluence of blue light (15 μmol/m²/s) while OsbZIP1.2 becomes dominant as the fluence is increased to 30 μmol/m²/s. Data presented in this study reveal that AtHY5 homologs in rice have undergone neofunctionalization and alternative splicing (AS) of *OsbZIP1* has increased the repertoire of its functions.

**One sentence summary:** Alternative spliced forms of *OsbZIP1*, an *AtHY5* homolog in rice, regulate seedling development in response to light and dark

## Introduction

Plants have evolved an extensive network to respond to light which is an important environmental cue responsible for regulating their growth and development. Light acts as a stimulus initiating different developmental pathways in plants enabling them to adapt to the changing environmental conditions. The mechanisms by which light brings about the desired phenotypic changes in plants are quite complex. The cascade of light signal transduction involves a number of components, which can be broadly grouped as photoreceptors, early signalling factors, central integrators and downstream effectors (Chory, 2010). The photoreceptors like phytochromes, cryptochromes, phototropins and zeitlupes perceive the light signals of varying wavelengths and thus are at the top in the hierarchy of the signal transduction pathway (Möglich et al., 2010; Mishra and Khurana, 2017; Paik et al., 2019). The signal is then transmitted to the early signalling factors like phytochrome interacting factors (PIF’s), HFR1 and LAF1, which further relay the signal to the downstream effectors like HY5 that are responsible for bringing about a dramatic alteration in gene expression by regulating the expression of a plethora of genes involved in photomorphogenesis (Chen and Chory, 2011; Xu, 2020). The central integrators belong to the COP/DET/FUS class of proteins which bring about the targeted degradation of other components involved in the light signal transduction pathway and thus integrate signals from all classes of photoreceptors, which regulate its activity in some way or the other (Hoecker, 2017; Bhatnagar et al., 2020; Xu, 2020).

*Arabidopsis* mutants exhibiting long hypocotyl in the presence of light as compared to the wild type have led to the characterization of HY5, a major downstream effector of the light signalling pathway (Koornneef et al., 1980; Oyama et al., 1997; Holm et al., 2002; Sibout et al., 2006; Gangappa and Botto 2016). The *HY5* gene encodes a nuclear localized basic leucine zipper transcription factor (168 aa) that consists of a N-terminal COP1 binding domain (25-60 aa) responsible for interaction with WD-40 domain of the COP1 dimer and a C-terminal bZIP domain (86-150 aa) which regulates DNA binding, homodimerization and heterodimerization (Bhatnagar et al., 2020). HY5 is phosphorylated at Ser36 within the COP1 binding domain by CASEIN KINASE 2 that reduces its susceptibility to degradation by COP1 (Hardtke, 2000). The unphosphorylated HY5 binds more efficiently to the promoters of light regulated genes like *CHS* (chalcone synthase) and *RBCS* (ribulose bis-phosphate carboxylase small subunit) (Ang et al., 1998; Chattopadhyay et al., 1998; Hardtke et al., 2000). *Arabidopsis* transgenic plants overexpressing full-length *HY5* do not show any phenotypic difference in hypocotyl length as compared to the wild type in light or dark except for a reduced lateral root density (van Gelderen et al., 2018). However, the overexpression of partial HY5 lacking the COP1 binding domain results in a hyper- photomorphogenic phenotype in *Arabidopsis* with the seedlings exhibiting a reduced hypocotyl length as well as anthocyanin levels under continuous red, far-red, blue and white light conditions with no deviant phenotype being observed in dark (Ang et al., 1998). HY5 acts as a central integrator for phytochromes, cryptochromes and UVR8, functioning as a positive regulator of photomorphogenesis under a broad spectrum of light (Ulm et al., 2004; Jiao et al., 2007; Brown and Jenkins 2008; Huang et al., 2012; Jiang et al., 2012; Singh et al., 2012; Wang et al., 2019). The degree of photomorphogenic development in *Arabidopsis* is directly proportional to the abundance of HY5 protein (Osterlund et al., 2000).

A HY5 homolog in *Arabidopsis,* HYH, has also been extensively characterized and shown to function in blue light mediated development and regulation of gene expression. Like HY5, it physically interacts with COP1 as a result of which it undergoes degradation in dark (Holm et al., 2002). It has also been reported to undergo alternative splicing (AS) induced by light (Li et al., 2017). Light regulates gene expression at every stage - transcriptional, translational, post-translational and probably pre-mRNA stage also (Liu et al., 2012). Light regulated alternative splicing can change the transcriptome globally and photoreceptors have been reported to participate in this process (Shikata et al., 2014; Wu et al., 2014; Cheng et al., 2018). Other than HYH, no other HY5 homolog has been reported to undergo alternative splicing in other species till now.

While HY5 is very well characterized in *Arabidopsis*, functions of its homologs are less understood in monocots. Subsequent to the availability of rice genome sequence (IRGSP, 2005), in which our group also participated, we carried out an extensive microarray analyses of the genes expressing differentially during reproductive development and abiotic stress in rice (Sharma et al., 2012). Using this microarray data generated in-house, our laboratory identified that the bZIP transcription family comprises of 89 members (Nijhawan et al., 2008). Among these, three homologs of AtHY5 in rice, namely *OsbZIP1*, *OsbZIP48* and *OsbZIP18*, were identified (Nijhawan et al., 2008; Burman et al., 2018). *OsbZIP48* has been shown to functionally complement the *Arabidopsis hy5* mutant. Interestingly, this *AtHY5* homolog in rice functions differently than its counterpart in *Arabidopsis*. While AtHY5 protein levels are light regulated, OsbZIP48 does not show light regulation at the protein level as it is not degraded in dark. Rather, it is developmentally regulated (Burman et al., 2018). The overexpression of *OsbZIP48* in rice leads to reduction in culm length, thus reducing plant height. OsbZIP48 has been shown to bind to the *OsKO2* (encoding ent- kaurene oxidase 2 enzyme of the gibberellin biosynthesis pathway) promoter thus, regulating plant height by affecting gibberellin biosynthesis pathway (Burman et al., 2018). Further investigation is necessary to find out if the AtHY5 homologs have acquired some new functions in other species as well. Recently, OsbZIP18, another AtHY5 homolog that we identified (Nijhawan et al., 2008) has been characterised by another group and they reported that OsbZIP18 upregulates the branched chain amino acids (BCAA) synthesis by binding to the ACE and C box elements in the promoters of *branched-chain aminotransferase1* Os*BCAT1* and *OsBCAT2* biosynthetic genes (Sun et al., 2020). The authors have shown that nitrogen deficiency causes induction of OsbZIP18, which in turn increases BCAA levels. Thus, it seems that OsbZIP18 has acquired a new function in monocots. *OsbZIP18* is also induced by UV-B stress and its overexpression in rice leads to a UV-B stress sensitive phenotype (Sun et al. 2022). In this study, we have functionally characterized OsbZIP1, another AtHY5 homolog in rice, to decipher if it has undergone neofunctionalization in rice. In addition, we have also explored the functional significance of alternative splicing of *OsbZIP1* in photomorphogenesis.

## Results

### Alternatively spliced transcripts of *OsbZIP1* exhibit developmental stage and light regulated expression

In order to elucidate the evolutionary divergence and neofunctionalization of *HY5* homologs in rice, we functionally characterized rice *OsbZIP1* in this study. We attempted to amplify the 612 bp coding sequence (CDS) of *OsbZIP1* (LOC_Os01g07880) using PCR from cDNAs prepared from various rice tissues as predicted by Rice Genome Annotation Project (Kawahara et al., 2013), however, a product of only 345 bp could be amplified. We performed 5’ Rapid Amplification of cDNA ends (RACE) followed by sequencing of the amplified products (345 bp and 612 bp) that confirmed an alternatively spliced form of *OsbZIP1* coding for a protein without the N-terminal COP1 binding domain in addition to a full-length protein having all the predicted functional domains (Fig. 1a). Analysis of the *OsbZIP1* gene structure revealed that the 780 bp genomic region of *OsbZIP1* can give rise to a 612 bp transcript consisting of three exons (hereafter referred as *OsbZIP1.1*) while the 345 bp alternatively spliced transcript retains only the partial regions of first and second exon and an intact third exon (hereafter referred as *OsbZIP1.2*) The full-length 612 bp *OsbZIP1.1* CDS possessing all the functional domains, as predicted by SMART (http://smart.embl-heidelberg.de/), was amplified by 5’ RACE using cDNA prepared from 10-day-old rice seedlings overexpressing *OsbZIP48* (OsbZIP48^OE^) as *OsbZIP1* was found to be upregulated (relative fold change = 9) in OsbZIP48^OE^ seedlings (Burman et. al., 2018).

**Figure 1.**
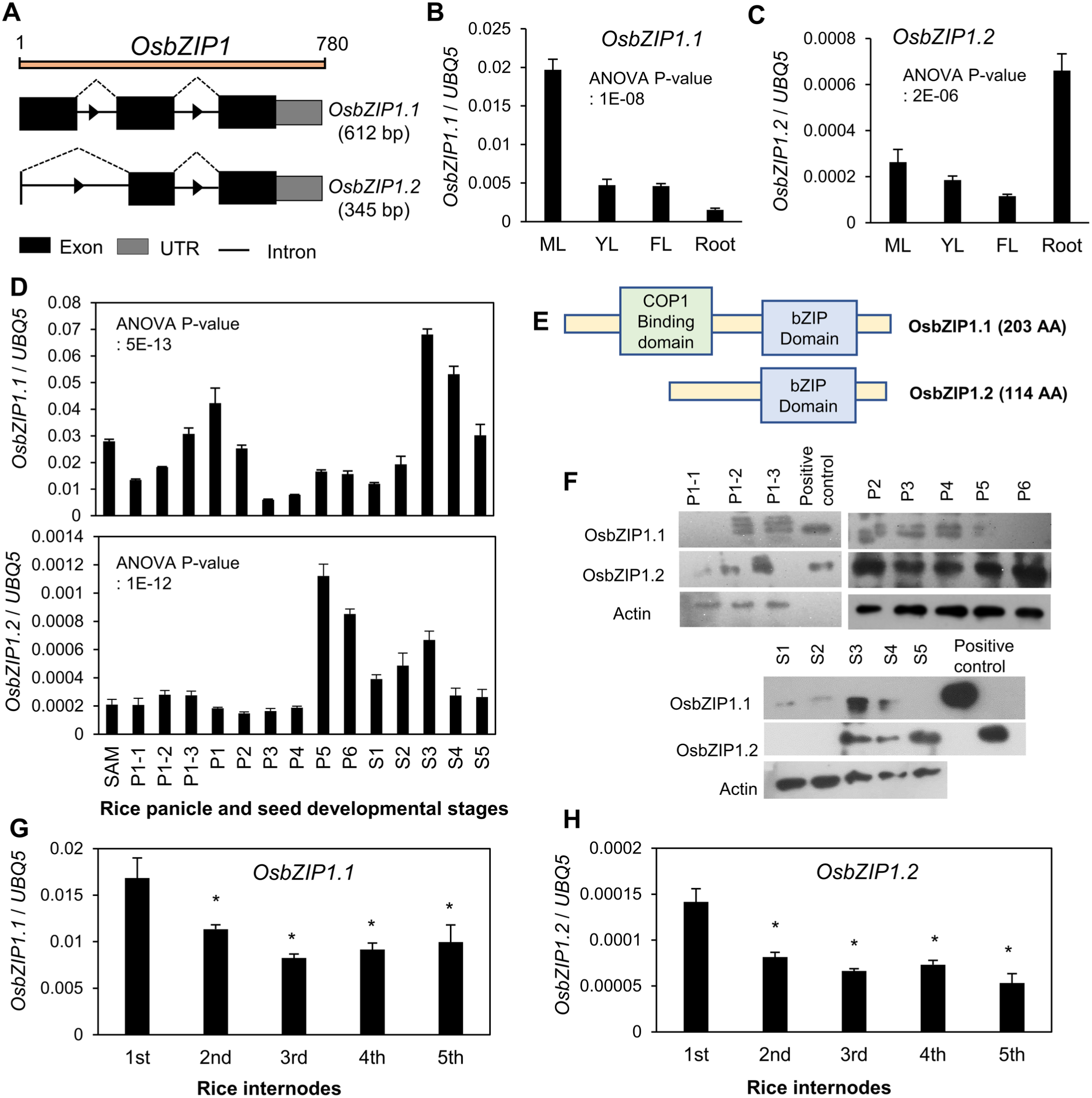
Expression profile of *OsbZIP1.1* and *OsbZIP1.2* in different tissues and developmental stages of rice. (**A)** Schematic diagram depicting the gene structure of *OsbZIP1.1* and *OsbZIP1.2*. Expression analysis of (**B)** *OsbZIP1.1* and **(C)** *OsbZIP1.2* in different vegetative tissues of rice [mature leaf (ML), Y-leaf (YL), flag leaf (FL) and root] carried out by RT-qPCR. ANOVA P-value < 0.05. **(D)** Expression of *OsbZIP1.1* and *OsbZIP1.2* in panicle and seed developmental stages of rice as analyzed by RT-qPCR (ANOVA P-value < 0.05). Details of stages are described in Material and Methods. **(E)** Schematic diagram depicting the domain organisation of OsbZIP1.1 and OsbZIP1.2. **(F)** Immunoblot showing protein levels of OsbZIP1.1 and OsbZIP1.2 in panicle development stages and in seed development stages of rice. Positive control is bacterially expressed and purified 6X-His-OsbZIP1.1/OsbZIP1.2. Anti- OsbZIP1 peptide antibody was used to immunoblot. Changes in transcript levels of **(G)** *OsbZIP1.1* and **(H)** *OsbZIP1.2* analysed by RT-qPCR in rice internodes. Data shown are means ± SD. The values shown for RT-qPCR experiments are relative transcript levels with respect to *UBIQUITIN5.* The diagrams in A and E are for representation only and not to scale.

The microarray-based genome wide expression profile (GSE6893 and GSE14298) showed that *OsbZIP1* transcript levels were maximum in root tissues, panicles (P2 stage of development) and in seeds (S5 stage of development) (Fig. S1A). Real-time quantitative PCR (RT-qPCR) analysis was carried out to validate the expression profile of *OsbZIP1*. The expression profiles of *OsbZIP1.1* and *OsbZIP1.2* were found to be slightly different from each other (Fig. 1B, C, D). While the levels of *OsbZIP1.1* transcript were high in mature leaf, P1 stage of panicle development, S3 and S4 stage of seed development; *OsbZIP1*.2 transcript levels were high at later stages of panicle development (P5 and P6). Further, *OsbZIP1.2* showed high expression in root as compared to the mature leaf and Y-leaf (Fig. 1C, S1B). This indicates that the *OsbZIP1* transcript levels are developmentally regulated and are likely tissue specific. The *in-vivo* protein profile of OsbZIP1.1 and OsbZIP1.2 was also studied in the panicle and seed development stages by western blotting using a custom-made peptide antibody that detected both the isoforms. A single band running at a molecular weight of ∼25 kDa was observed for OsbZIP1.2, while 2-3 bands running at a molecular weight of ∼33- 35 kDa were observed for OsbZIP1.1 in different tissues. This may be due to differentially phosphorylated forms of OsbZIP1.1 as has been reported for other light signaling components earlier (Kim et al., 2004; Shin et al., 2016; Hoang et al., 2019). We observed sustained OsbZIP1.1 level during P1-2 to P4 stage after which it declined at later stages of the panicle development. OsbZIP1.2 protein was observed from P1-2 stage after which it remained stable up to P6 stage. During seed developmental stages, OsbZIP1.1 was observed at higher levels in the S3 stage while OsbZIP1.2 was observed in S3, S4 and S5 stages of seed development (Fig. 1F). Overall, the transcript profile of OsbZIP1.1 and OsbZIP1.2 was essentially similar to their protein profile. The expression pattern of *OsbZIP1.1* and *OsbZIP1.2* was also similar in the internodes of rice with both of them expressing maximum in the first internode of rice and declining thereafter (Fig. 1G, H).

To monitor the effect of light/dark conditions on the expression of *OsbZIP1*, we first studied the seedling development in dark. The 3-day-old dark-grown rice seedling consists of a coleoptile and root. The first (primary) leaf is enclosed within the coleoptile in a 4-day-old seedling after which it emerges out of the coleoptile on day 5 (Fig. 2A). The second leaf sheath is observed on day 6, after which it elongates till day 7 and the second leaf blade emerges by day 8. It continues to elongate till day 10. We monitored the transcript levels of *OsbZIP1.1* as well as *OsbZIP1.2* using RT-qPCR every 24 hours from day 2 onwards till day 10 in the light and dark grown rice seedlings (Fig. 2B, C). The expression of *OsbZIP1.1* was found to be maximum in the 2-day-old light-grown seedlings while declining gradually from day 3 onwards till day 10. However, the expression of *OsbZIP1.2* increased from day 2 till day 4 and decreased thereafter in light-grown seedlings. (Fig. 2B, C). In dark-grown seedlings, the expression of *OsbZIP1.2* was particularly high at 4^th^ day and 7^th^ day, around the time when the first and second leaves emerge. We observed its maximum expression in 10-day-old dark grown seedlings (Fig. 2C). This indicates a role of *OsbZIP1.2* in leaf emergence and development under dark conditions. The expression of *OsbZIP1.1* increases gradually in 2- to 4-day-old dark-grown seedlings, decreases thereafter and peaks again on 9^th^ day (Fig. 2B).

**Figure 2.**
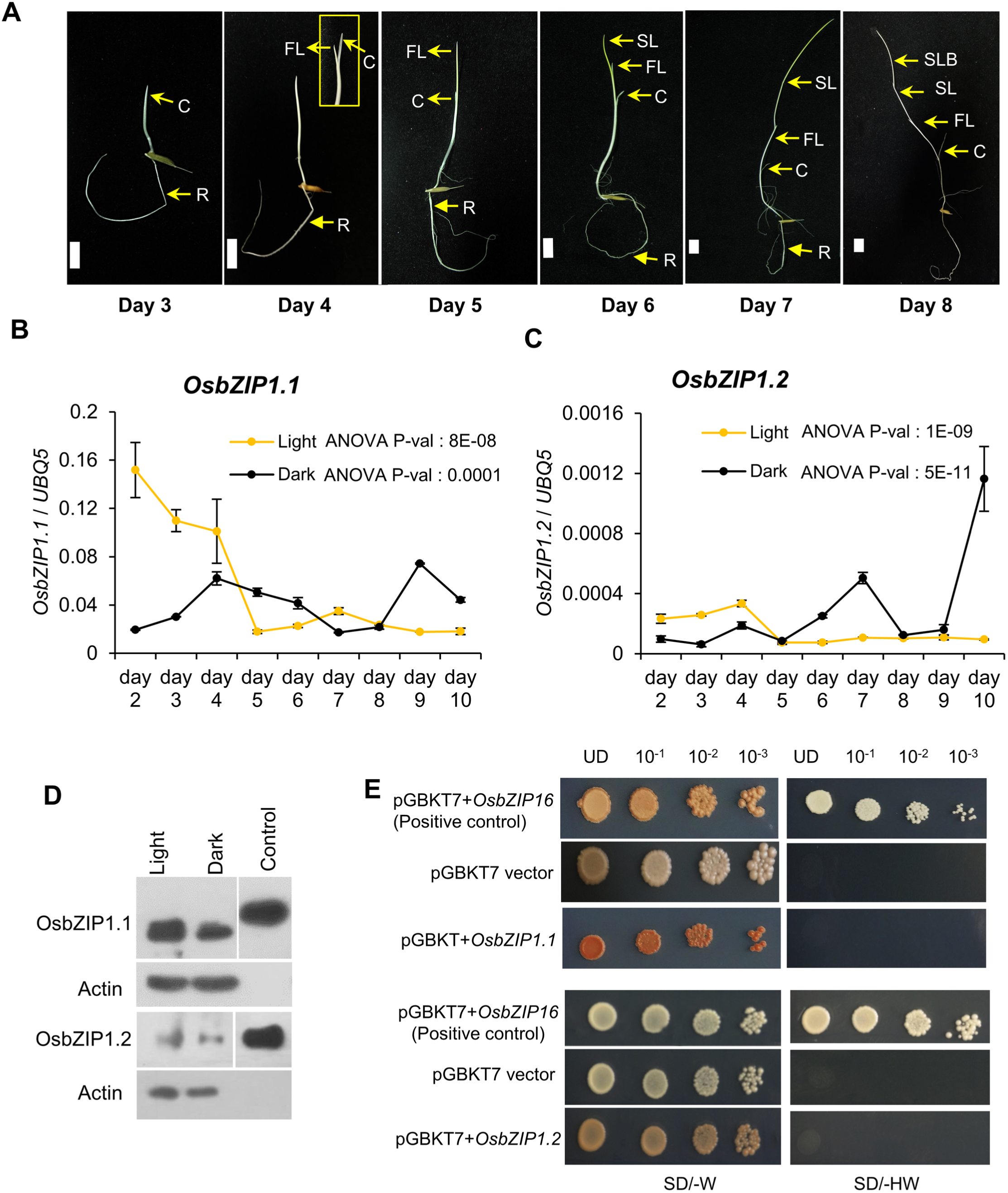
Light/dark-regulated levels of OsbZIP1.1 and OsbZIP1.2. (A) Development of rice seedling in dark showing the emergence of first and second leaves. C, coleoptile; R, root; FL, first leaf; SL, second leaf; SLB, second leaf blade. Relative expression of *OsbZIP1.1* and *OsbZIP1.2* in 2-to-10-day-old **(B)** light-grown **(C)** dark-grown seedlings of rice as analysed by RT-qPCR. Data shown are means ± SD, ANOVA P-value < 0.05. The expression data presented are relative to *UBIQUITIN5*. **(D)** Immunoblot showing the levels of OsbZIP1.1 and OsbZIP1.2 in 7-day-old light- and dark-grown calli overexpressing the genomic region of *OsbZIP1*. The positive control is bacterially expressed and purified 6X-His-OsbZIP1.1/OsbZIP1.2 which were run on the same gel as the other samples. **(E)** Transactivation assay of OsbZIP1.1 and OsbZIP1.2 in yeast. UD (undiluted), 10−^1^, 10−², 10−^3^ represent the dilutions of the yeast cultures used. SD/-HW (medium without histidine and tryptophan); SD/-W (medium without tryptophan).

To ascertain the effect of only light/dark and not the developmental stage of rice on levels of OsbZIP1 protein isoforms, we checked the 7-day-old light- and dark-grown calli obtained from rice seeds overexpressing *OsbZIP1* genomic DNA (gDNA). The protein levels of OsbZIP1.1 were higher in light as compared to dark while that of OsbZIP1.2 were similar under both the conditions, implying that the protein levels of *OsbZIP1.1* are regulated by light (Fig. 2D).

### OsbZIP1.1 and OsbZIP1.2 lack transactivation ability, show differential subcellular localization and can form homo- or hetero-dimers

Some bZIP transcription factors like OsbZIP16 are known to have a transactivation domain due to which they are able to activate the transcription of downstream genes (Chen et al., 2012; Pandey et al., 2018). Thus, the transactivation ability of OsbZIP1.1 and OsbZIP1.2 was checked in yeast. The yeast cells expressing either *OsbZIP1.1* or *OsbZIP1.2* were unable to grow on SD/-HW medium implying that both OsbZIP1.1 and OsbZIP1.2 lack the transactivation activity (Fig. 2E). OsbZIP16 was used as the positive control in this experiment and it was able to grow on SD/-HW medium while empty pGBKT7 vector was used as the negative control.

To identify the sub-cellular localization of OsbZIP1.1 and OsbZIP1.2, particle bombardment was carried out in onion epidermal cells. The YFP-OsbZIP1.1 fusion protein was found to be localized in the nucleus while YFP-OsbZIP1.2 was found to localize all over the cell including nucleus (Fig. 3A). Since, the bZIP transcription factors consist of a basic leucine zipper domain which is involved in the dimerization of two proteins (Landschulz et al., 1998), we checked homodimerization of OsbZIP1.1 *in vivo* by Bimolecular Fluorescence complementation (BiFC) assay using split YFP system. *OsbZIP1.1* fused separately to N-terminal YFP and C-terminal YFP were co-bombarded in onion epidermal cells and fluorescence indicating physical interaction was detected in the nucleus (Fig. 3B). This was further validated by yeast 2-hybrid (Y2H) assay using co-transformants (pGBKT7-*OsbZIP1.1* and pGADT7-*OsbZIP1.1*) grown on SD/-HLW medium while no growth was observed in case of the negative control (pGBKT7-*Lam* and pGADT7-*T*), implying that the full-length OsbZIP1 (OsbZIP1.1) can homodimerize (Fig. 3C). Similarly, OsbZIP1.2 was also checked for homodimerization by BiFC as well as by Y2H that confirmed positive interaction among OsbZIP1.2 protein monomers. Thus, both OsbZIP1.1 and OsbZIP1.2 can homodimerize in the nucleus.

**Figure 3.**
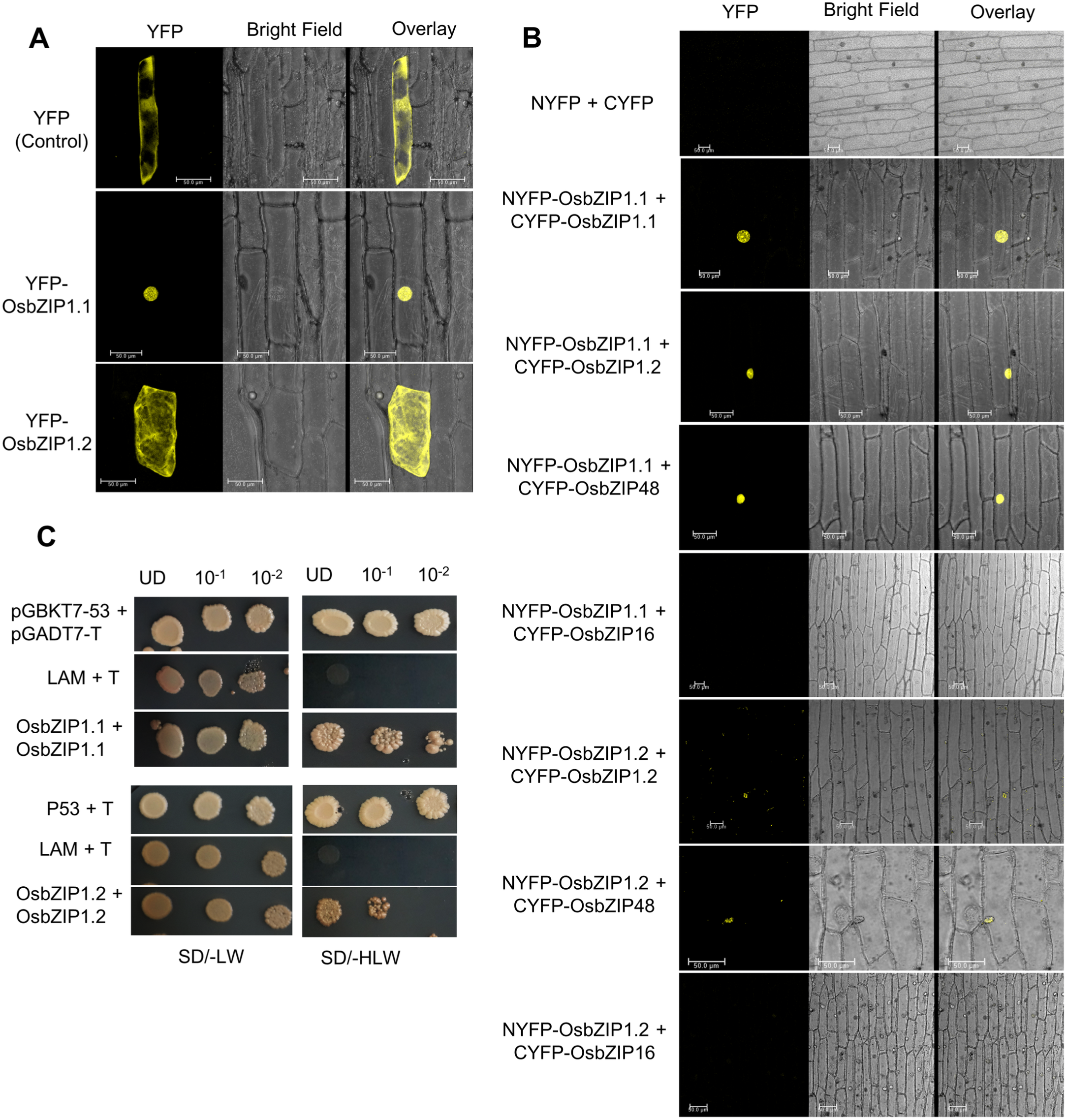
Subcellular localisation, homo and heterodimerization of OsbZIP1.1 and OsbZIP1.2. (A) Sub-cellular localization of OsbZIP1.1 and OsbZIP1.2 in onion peel cells. The first row (YFP control vector) shows localization of only YFP protein, the second and third rows show localization of YFP-OsbZIP1.1 and YFP-OsbZIP1.2, respectively. **(B)** Bimolecular Fluorescence Complementation (BiFC) analysis in onion peel cells showing the homodimerization of OsbZIP1.1 and OsbZIP1.2, heterodimerization of OsbZIP1.1 with OsbZIP1.2 and that of OsbZIP1.1 with OsbZIP48 as well as OsbZIP1.2 with OsbZIP48 in the nucleus. OsbZIP16 is used as a negative control. The first column shows photographs taken using YFP filter, and the second column shows bright field image of the same cell, the third column depicts the merged photographs of bright field and fluorescence images taken using a Leica microscope. **(C)** Yeast-two-hybrid assay showing homodimerization of OsbZIP1.1 and OsbZIP1.2. UD (undiluted), 10−^1^, 10−² represent the dilutions of the yeast cultures used. SD/-LW (medium without Leucine and tryptophan); SD/-HLW (medium without histidine, leucine and tryptophan).

Previously on the basis of predicted dimerization specificities, we divided the bZIP transcription factor family members into 29 subfamilies (BZ1-BZ29) (Nijhawan et al., 2008). OsbZIP1 (LOC_Os01g07880), OsbZIP48 (LOC_Os06g39960) and OsbZIP18 (LOC_Os02g10860) were grouped together as BZ2 and have been predicted to homodimerize as well as heterodimerize with the same group members. Since, both OsbZIP48 and OsbZIP1.1 show nuclear localization upon transient expression, we checked if OsbZIP1.1 and OsbZIP1.2 can heterodimerize with OsbZIP48. We observed physical interaction between the two using BiFC assays (Fig. 3B). Fluorescence was observed in the nucleus in both the cases implying that both OsbZIP1.1 and OsbZIP1.2 heterodimerize with another HY5 homolog OsbZIP48. However, both OsbZIP1.1 and OsbZIP1.2 could not heterodimerize with another bZIP transcription factor OsbZIP16 indicating the specialized nature of this interaction among bZIP family members. Thus, OsbZIP1.1 can form homodimers with itself and heterodimers with OsbZIP1.2 and OsbZIP48.

### OsbZIP1.1 and OsbZIP1.2 can complement Arabidopsis hy5 mutant

*Arabidopsis hy5* mutants are known to exhibit an elongated hypocotyl, reduced greening especially in the middle and lower parts of the hypocotyl, reduced cotyledon angle, increased number of lateral roots (LR) and LR angle when grown under white light (Srivastava et al., 2015; Ang et al., 1998, Oyama et al., 1997; Chattopadhyay et al., 1998, Burman et al., 2018, van Gelderen et al., 2018). We confirmed that *OsbZIP1* is indeed a functional homolog of *AtHY5* by complementation of *Arabidopsis hy5* mutant with *OsbZIP1 (gDNA)* and *OsbZIP1.2*. The complementation lines thus, raised were checked for the presence of hygromycin resistance marker gene by PCR (Fig. S2B, D).

The hypocotyl length of 6-day-old complementation *hy5*/35S:*OsbZIP1*, *hy5*/35S:*OsbZIP1.2* as well as overexpression WT/35S:*OsbZIP1* and WT/35S:*OsbZIP1.2* lines were similar to the wild type under white light conditions (Fig. 4A, B, E, S3A, S3B). Further, the chlorophyll content of overexpression as well as the complementation lines of *OsbZIP1* and *OsbZIP1.2* was also similar to the wild type and significantly more than the *hy5* mutant (Fig. S3C, S4A). Overexpression of either *OsbZIP1* or *OsbZIP1.2* also showed an average cotyledon angle, lateral roots phenotype similar to the wild type (Figure 4B, 4C, S5).

**Figure 4.**
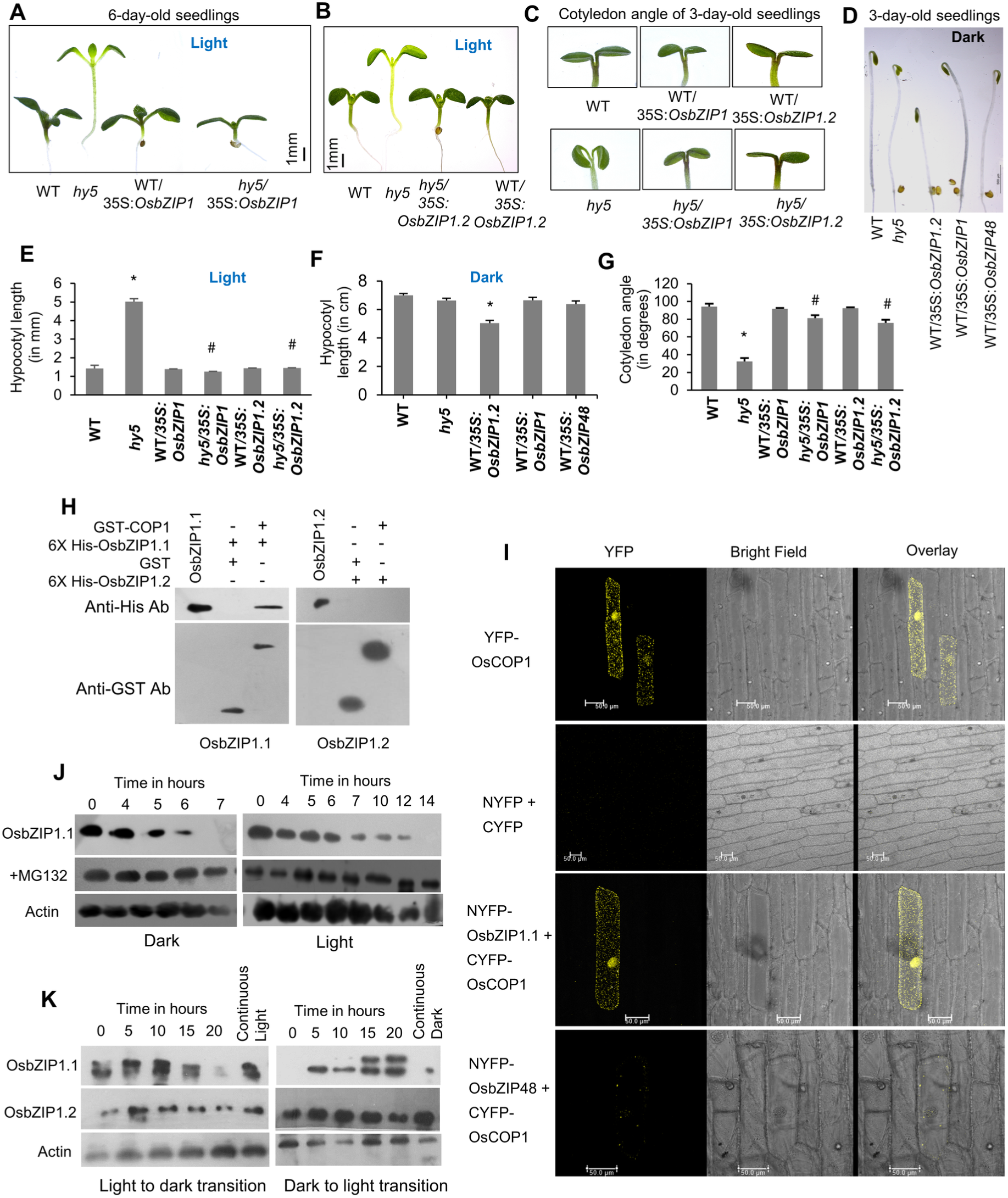
Complementation of *Athy5* mutant by *OsbZIP1* and its interaction with OsCOP1. (A) Representative picture shows the phenotype of 6-day-old white light-grown wild-type (WT), *hy5*, WT/35S:*OsbZIP1*, and *hy5*/35S:*OsbZIP1* seedlings and **(B)** WT/35S:*OsbZIP1.2* and *hy5*/35S:*OsbZIP1.2* seedlings. **(C)** Representative picture shows cotyledon opening angle of 3-day-old white light-grown seedlings. **(D)** Representative picture shows the phenotype of 3- day-old dark-grown seedlings. **(E)** Histograms show hypocotyl lengths of 6-day-old white light-grown seedlings and **(F)** hypocotyl lengths of 3-day-old dark-grown seedlings. **(G)** Histograms show measurements of cotyledon opening angle of 3-day-old white light-grown seedlings. T-test p-value < 0.01 {Asterisk (*) shows comparison with WT and hash (#) with *hy5* mutant}. Data shown is mean ±SE, n=30 seedlings **(H)** *In vitro* GST-pull down assay showing interaction of OsbZIP1.1 with OsCOP1 and non-interaction of OsbZIP1.2 with OsCOP1. **(I**) First row shows the sub-cellular localisation of OsCOP1 in onion epidermal cells, second row shows the Bimolecular Fluorescence Complementation (BiFC) assay using empty vectors as negative control, third row shows the BiFC assay in which OsbZIP1.1 interacts with OsCOP1 and fourth row shows non-interaction of OsbZIP48 with OsCOP1. OsbZIP48 acts as a negative control. **(J)** *In vitro* degradation assay showing degradation of 6X-His- OsbZIP1.1 within 7 h when incubated with protein extract of dark-grown seedlings and within 14 h when incubated with protein extract of light-grown seedlings for the specified time points. MG132 was used as the inhibitor of 26S proteasome. Actin was used as the loading control. Anti-His antibody was used to immunoblot the levels of OsbZIP1.1. **(K)** *In vivo* degradation of OsbZIP1.1 within 20 hrs of transfer of 4-day-old light grown seedlings to dark and increase in OsbZIP1.1 protein levels after transfer of 4-day-old dark grown seedlings to light within 5 hrs whereas no drastic change in the protein levels of OsbZIP1.2 was observed upon light to dark and dark to light transition. Control used in light to dark transition or dark to light transition experiment was 4-day-old continuous light- or dark- grown rice seedlings, respectively. Anti-OsbZIP1 peptide antibody was used to immunoblot levels of both the isoforms. Actin was used as loading control.

Over-expression of full length *AtHY5* in *Arabidopsis* is unable to cause any change in hypocotyl length of seedlings in white light (Ang et al., 1998). However, over-expression of AtHY5-NΔ77 (77 aa deletion at N-terminal) with a deletion of COP1 binding domain leads to a hyper-photomorphogenic phenotype in *Arabidopsis* with enhanced reduction in hypocotyl length under white light as well as monochromatic lights (red, blue and far-red) (Ang et al., 1998). Thus, we overexpressed *OsbZIP1.2* that lacks the COP1 binding domain in *Arabidopsis* and observed that the average hypocotyl length of these transgenics was similar to the wild type under white light unlike AtHY5-NΔ77 seedlings (Fig. 4B, E). Interestingly, the average hypocotyl length of WT/35S:*OsbZIP1.2* seedlings was significantly reduced under dark conditions while that of WT/35S:OsbZIP1 and WT/35S:OsbZIP48 was similar to the wild type seedlings (Fig. 4D, 4F). However, opening of apical hook, a characteristic feature of photomorphogenic development was not observed under dark conditions. Thus, overexpression of *OsbZIP1.2* lacking a COP1 binding domain causes reduction in hypocotyl length of 3-day-old dark-grown *Arabidopsis* seedlings, which is not observed in case of ectopic expression of *AtHY5-NΔ77* in *Arabidopsis* seedlings grown in dark (Ang et al., 1998). Therefore, while OsbZIP1 and *OsbZIP1.2* are able to complement the *hy5* Arabidopsis mutant phenotype in terms of hypocotyl length, cotyledon angle and chlorophyll under light, overexpression of *OsbZIP1.2* leads to a partial photomorphogenic response in dark.

### OsbZIP1.1 is targeted by OsCOP1

In order to decipher if OsbZIP1 works in the same way as AtHY5, we checked the interaction of OsbZIP1.1 and OsbZIP1.2 with OsCOP1 using *in vitro* GST pull-down assays that showed OsCOP1 can interact with OsbZIP1.1 but not with OsbZIP1.2 (Fig. 4H). The interaction of OsbZIP1.1 with OsCOP1 was further observed using BiFC as bright speckles all over the cell as well as in the nucleus of onion epidermal cells (Figure 4I). While YFP-OsbZIP1.1 is localized in the nucleus *in vivo*, YFP-OsCOP1 localization was observed as a speckled distribution in the nucleus and all over the cell. This interaction between OsbZIP1.1-OsCOP1 is essentially similar to AtHY5-AtCOP1 interaction reported earlier by Ang et al. (1998). We did not observe any interaction of OsCOP1 with OsbZIP48, another *AtHY5* homolog in rice (Fig. 4I, S6A).

OsCOP1, an E3 ubiquitin ligase, is known for degradation of a number of photomorphogenesis promoting transcription factors (von Arnim et al., 1997; Holm et al., 2002; Seo et al., 2003; Jang et al., 2005; Lin et al., 2018; Bhatnagar et al., 2020). *In vitro* degradation assay performed using cell free system with 6X-His-OsbZIP1.1 showed that OsbZIP1.1 got degraded within 7 hours using dark-grown seedlings extract, the same amount of OsbZIP1 got degraded within 14 hours when light-grown seedlings extract was used (Fig. 4J). No degradation of the OsbZIP1.1 protein was observed when MG132, an inhibitor of 26S proteasome, was added to the reaction mixture, suggesting that the degradation of OsbZIP1.1 is mediated by the 26S proteasome pathway. *In vivo* degradation of OsbZIP1.1 was also examined by subjecting the 4-day-old light-grown rice seedlings to darkness for 5,10, 15 and 20 hours and analyzing the protein levels of OsbZIP1.1 and OsbZIP1.2 by immunoblotting. OsbZIP1.1 was found to degrade within 20 hours of transfer of the 4-day-old rice seedlings from light to dark whereas the OsbZIP1.2 levels remained largely unchanged. This is in consonance with observations mentioned above that only OsbZIP1.1 appears to be light regulated. We further checked if any changes occur in the protein levels of OsbZIP1.1 and OsbZIP1.2 by immunoblotting during dark to light transition by subjecting the 4-day-old dark grown rice seedlings to light for 5, 10, 15 and 20 hours. The unphosphorylated form of OsbZIP1.1 was observed within 5 hour of incubation in light and the phosphorylated form of OsbZIP1.1 appeared on the immunoblot within 10-15 hours as the levels of OsbZIP1.1 increased upon dark to light transition, indicating the light regulation of OsbZIP1.1. The levels of OsbZIP1.2 were almost similar during dark to light transition (Fig. 4K).

### OsbZIP1 interacts with OsCASEIN KINASE 2 and undergoes phosphorylation

Since many components of the light signal transduction pathway are known to be regulated by phosphorylation (Kim et al., 2004; Shin et al., 2016; Hoang et al., 2019) and we observed the presence of double bands whenever OsbZIP1.1 protein was subjected to immunoblotting, we studied whether OsbZIP1.1 shows a post-translational modification by phosphorylation. Since, AtHY5 was shown to undergo phosphorylation at the conserved CASEIN KINASE 2 (CK2) phosphorylation site located within the N-terminal COP1 binding domain (Hardtke et al., 2000), we analyzed the amino acid sequence of OsbZIP1.1. Interestingly, a conserved CK2 phosphorylation site was found to be present from amino acid 43 to 47 (ESDEE). Thus, the interaction of OsbZIP1.1 was checked with rice CK2. The four CK2 subunits (ɑ2, ɑ3, ß1 and ß3) were checked for their putative interaction with OsbZIP1.1 using Y2H. We found a positive interaction between OsbZIP1.1 and CK2ß1 as well as between OsbZIP1.1 and CK2ɑ3 (Fig. S6B). Since CK2ɑ3 is known to act as a catalytic subunit of CK2 (Filhol et al., 2004; Chen et al., 2015), we further confirmed the interaction of OsbZIP1 with CK2ɑ3 using BiFC. YFP-CK2ɑ3 was found to be localized all over the cell being predominantly present both in the nucleus as well as the cytoplasm (Fig. 5A). While cYFP- CKɑ3 was able to interact with nYFP-OsbZIP1.1 within the nucleus of cell, it showed no interaction with OsbZIP48 (Fig. 5A, S7). The interaction of OsbZIP1.1 with CK2ɑ3 was further confirmed by *in-vitro* GST pull-down assay (Fig. 5B). OsbZIP1.2 did not show any interaction with OsCK2ɑ3 (Fig. 5B).

**Figure 5.**
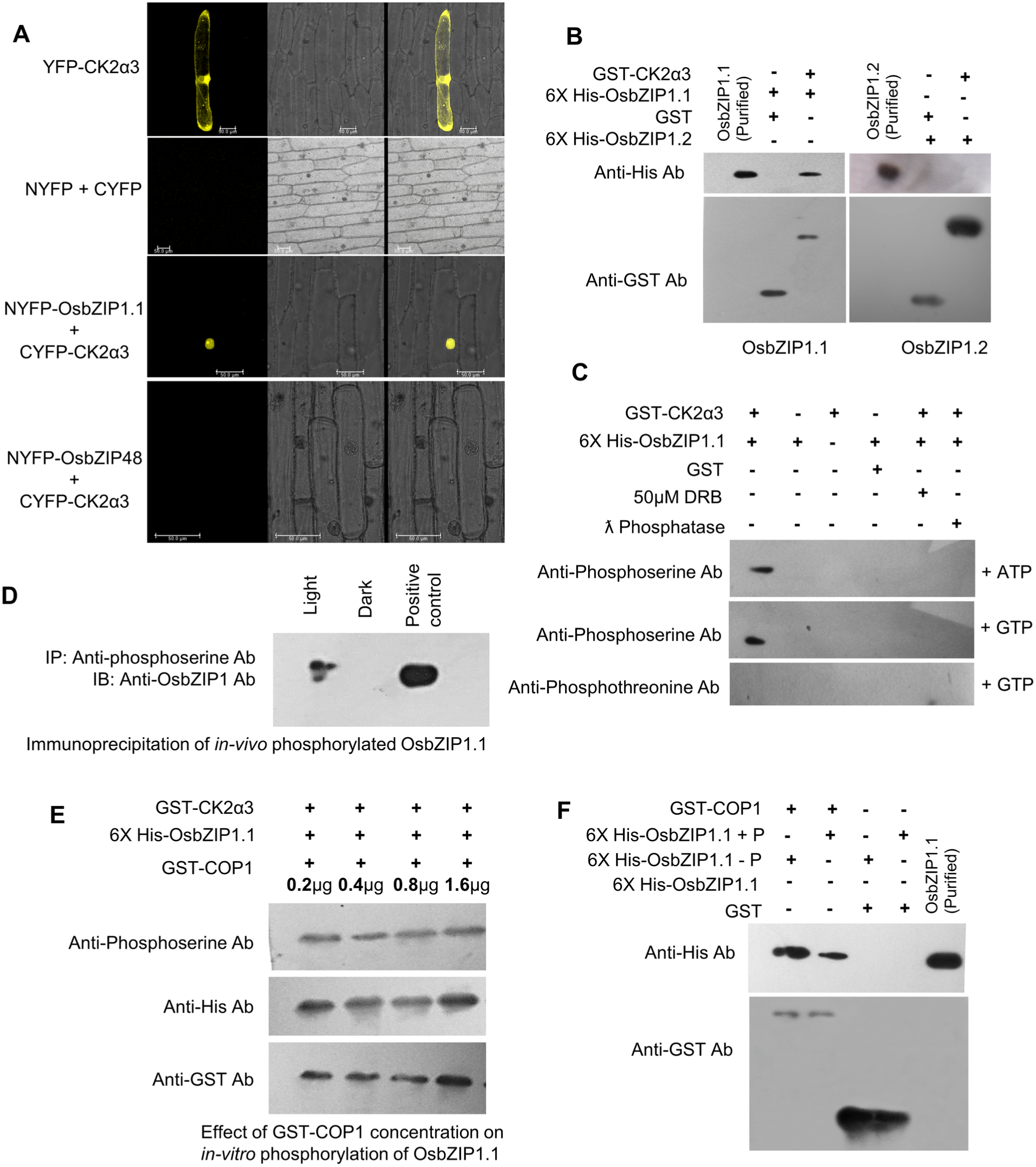
Interaction of OsbZIP1.1 with OsCK2ɑ3 and its subsequent phosphorylation. (A) First row shows subcellular localisation of OsCK2ɑ3, second row shows Bimolecular Fluorescence Complementation (BiFC) assay using empty vectors as negative control, third row shows the BiFC assay in which OsbZIP1.1 interacts with OsCK2ɑ3 in the nucleus of the onion epidermal cells and fourth row shows OsbZIP48 as the negative control showing non- interaction with OsCK2ɑ3. **(B)** *In vitro* GST-pull down assay showing interaction of OsbZIP1.1 with OsCK2ɑ3 and non-interaction of OsbZIP1.2 with OsCK2ɑ3. **(C)** *In vitro* kinase assay showing the phosphorylation of 6X-His-OsbZIP1.1 upon addition of GST-OsCK2ɑ3 and ATP or GTP. The blot was probed by anti-phosphoserine and anti-phosphothreonine antibodies. 50 µm DRB was used as a specific inhibitor of OsCK2ɑ3 and Lambda (ƛ) phosphatase was used as a non-specific inhibitor. GST was used as a negative control. **(D)** Immunoblot probed with anti-OsbZIP1 antibody showing the band of phosphorylated OsbZIP1.1 immunoprecipitated using anti-phosphoserine antibody from 5-day-old light-grown seedlings. No band was visible after immunoprecipitation from 5-day-old dark-grown seedlings. **(E)** Immunoblot showing the addition of increasing concentrations of GST-COP1 which does not affect the phosphorylation of 6X-His-OsbZIP1.1 in the *in vitro* phosphorylation assay. The blot was probed with anti-phosphoserine antibody and also with anti-His and anti-GST antibodies as control. **(F)** *In vitro* GST pull down assay showing interaction of *in vitro* phosphorylated and de-phosphorylated 6X-His-OsbZIP1 with GST-COP1. Bacterially expressed and purified GST protein was used as the negative control while the purified 6X-His-OsbZIP1.1 was used as the positive control. The blot was probed with anti-His antibody and also with anti-GST antibody as a control.

CASEIN KINASES (CK) are known to use GTP as an alternative phosphate donor (Hardtke et al., 2000). Thus, *in vitro* kinase assay carried out in the presence of either GTP or ATP as a phosphate donor showed that CK2ɑ3 is able to phosphorylate OsbZIP1.1 (Fig. 5C). The phosphorylation was found to take place at serine residues of OsbZIP1.1 as a band was detected using anti-phosphoserine antibody and not on the threonine residues as no band was observed when the membrane was probed with anti-phosphothreonine antibody (Fig. 5C). Notably, OsCK2ɑ3 did not show autophosphorylation activity (Fig. 5C). The specificity of the phosphorylation reaction was also confirmed by using 5,6-dichloro-1-beta-D-ribofuranosylbenzimidazole (DRB), a specific inhibitor of CASEIN KINASE 2 (CK2) (Kubinski et al., 2017). We further checked whether phosphorylation of OsbZIP1.1 by GST-CK2ɑ3 can be reversed with ƛ phosphatase, a phosphatase enzyme having affinity towards phosphorylated residues. Since no band was observed when either DRB or ƛ phosphatase was used to inhibit/reverse the activity of CK2 shows that CK2 is involved in the phosphorylation of OsbZIP1.1 (Fig 5C). Interestingly, OsbZIP48 exists in a phosphorylated state *in vivo* as a band is observed using anti-OsbZIP48 antibody when OsbZIP48 is immunoprecipitated using anti-phosphoserine antibody from 5-day- and 7-day-old light grown seedling extract (Fig. S7A). However, OsbZIP48 did not show interaction with OsCK2α3 (Fig. S7B).

The *in vivo* phosphorylation status of OsbZIP1.1 was also checked by immunoprecipitation of anti-phosphoserine proteins using anti-phosphoserine antibody from the 5-day-old light- and dark-grown seedling protein extract and subsequent immunoblotting using anti-OsbZIP1 antibody. A band obtained after immunoblotting ascertained the existence of OsbZIP1.1 in a phosphorylated state *in vivo* under light (Fig. 5D). As stated above, a conserved CK2 phosphorylation site (amino acid 43 to 47) is located within the COP1 binding domain of OsbZIP1.1 with the ser44 residue the most likely target of CK2 phosphorylation with a score of 0.723 as predicted by the NetPhos 3.1 server (http://www.cbs.dtu.dk/services/NetPhos/) (Fig. S8A). This was confirmed using mutated osbzip1-S44A (with serine at 44^th^ residue substituted with an alanine) where no phosphorylation by CK2 was detected using anti-phosphoserine antibody (Figure S8B).

Since OsCOP1 physically interacts with OsbZIP1.1, we checked whether OsCOP1 is able to affect the phosphorylation status of OsbZIP1.1. We observed that with increasing amount of GST-COP1 added to the reaction mixture containing 6X-His-OsbZIP1.1 and GST- CK2ɑ3 in the presence of GTP, the intensity of the single band obtained after western blotting, as detected by anti-phosphoserine antibody, remains the same (Fig. 5E). It implies that OsCOP1 does not interfere with the phosphorylation of OsbZIP1.1 by CK2ɑ3. We further detected that the unphosphorylated OsbZIP1.1 interacts with OsCOP1 more strongly than the phosphorylated OsbZIP1.1 *in vitro* (Fig. 5F). This was further confirmed with mutated osbzip1.1-S44A that interacts more strongly with OsCOP1 than the phosphorylated OsbZIP1.1 (Fig. S8C). This indicates that the unphosphorylated form of OsbZIP1.1 has a greater affinity for OsCOP1 and thus, it is more susceptible to degradation by OsCOP1 as compared to the phosphorylated form. In other words, phosphorylation protects OsbZIP1.1 from being degraded due to its lesser interaction or affinity with OsCOP1.

### Changes in OsbZIP1 levels in rice alters sensitivity to blue, red and far-red light

To functionally characterize *OsbZIP1* in rice, stable rice transgenics were raised for either overexpression of 780 bp genomic region of *OsbZIP1* (*OsbZIP1*^OE^) which led to overexpression of both the forms (*OsbZIP1.1* and *OsbZIP1.2*) or down-regulated expression of both spliced forms in RNAi (*OsbZIP1*^KD^) plants using *Agrobacterium*-mediated rice transformation (Fig.6A, S9 A). Rice transgenics with overexpression of only *OsbZIP1.2* were semi-dwarf and did not set seeds repeatedly. Therefore, we could not proceed with *OsbZIP1.2^OE^* and perform individual characterization of *OsbZIP1.1* and *OsbZIP1.2* in rice. Nevertheless, for *OsbZIP1*^OE^ and *OsbZIP1*^KD^, at least 10 transgenic lines were obtained and further work was extended on 3 lines for morphometric analyses. These lines were checked for the presence of hygromycin resistance gene by PCR (Fig. S9 B, C)

**Figure 6.**
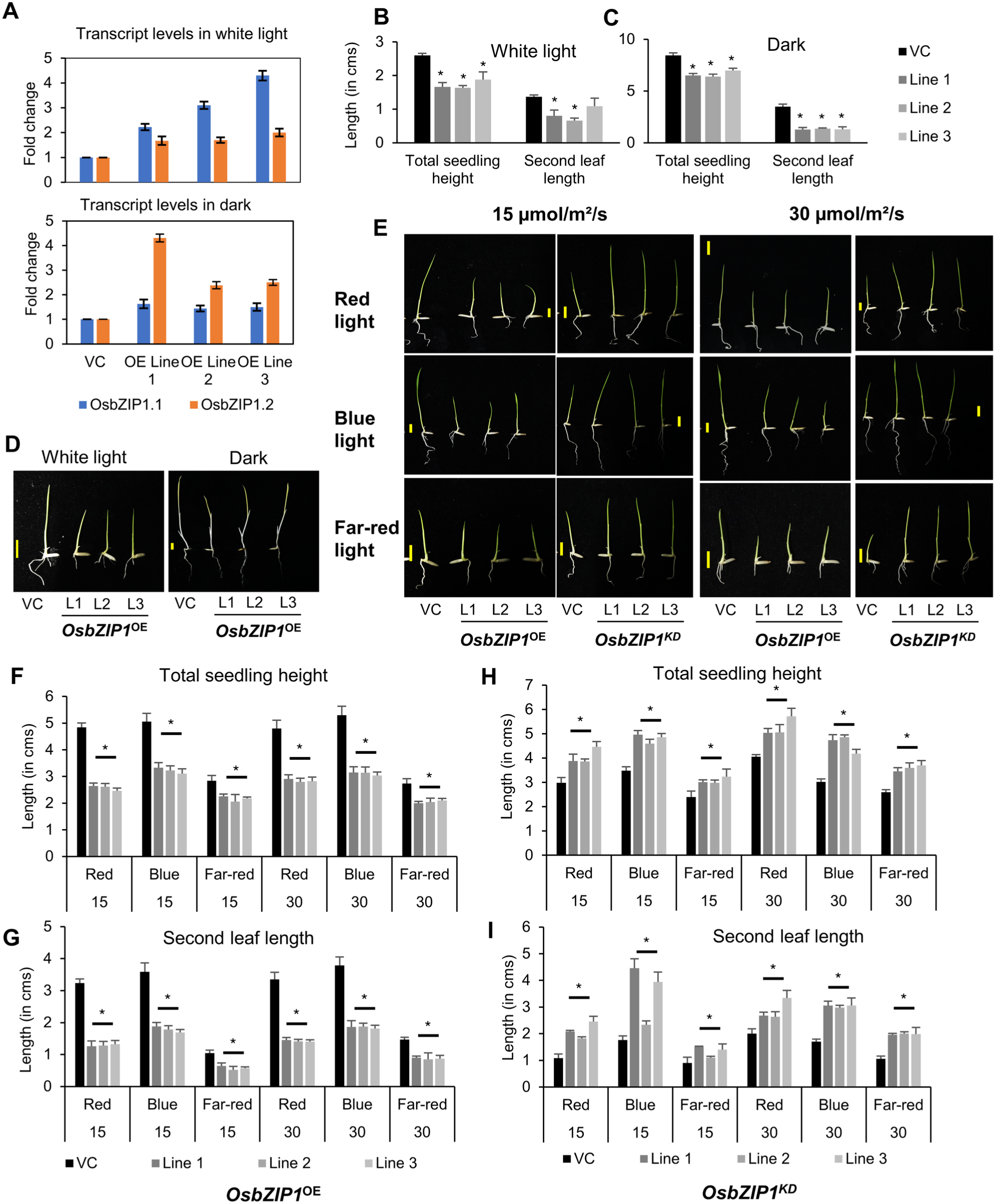
*OsbZIP1* regulates seedling height under monochromatic (white light, dark, red, far-red and blue) light conditions. (A) Fold change in transcript levels of *OsbZIP1.1* and *OsbZIP1.2* in the 5-day-old three transgenic lines of *OsbZIP1*^OE^ grown under continuous white light and dark conditions detected by RT-qPCR. **(B)** Total seedling height of 5-day-old *OsbZIP1*^OE^ grown in 100 µmol/m^2^/s of continuous white light and **(C)** in dark conditions is shown in the respective histograms. **(D)** Representative picture shows the phenotypes of 5- day-old *OsbZIP1*^OE^ seedlings grown in 100 µmol/m^2^/s of continuous white light or continuous dark. **(E**) Representative pictures show phenotypes of 5-day-old *OsbZIP1*^OE^ and *OsbZIP1*^KD^ rice seedlings grown under 15 and 30 µmol/m^2^/s of red, far-red or blue light conditions. **(F)** Total seedling height and **(G)** second leaf length (in cm) of 5-day-old *OsbZIP1*^OE^ rice seedlings grown under 15 and 30 µmol/m^2^/s of red, far-red and blue light. **(H)** Total seedling height and **(I)** second leaf length (in cm) of 5-day-old *OsbZIP1*^KD^ rice seedlings grown under 15 and 30 µmol/m^2^/s of red, far-red and blue light. Significance of value by T- test p-value < 0.01 in overexpression/RNAi plants vs respective vector control shown by asterisk (*).

When the transcript levels of the *OsbZIP1.1* and *OsbZIP1.2* were checked by RT-qPCR in the 5-d-old seedlings overexpressing *OsbZIP1* (gDNA), the *OsbZIP1.2* expression levels were higher than *OsbZIP1.1* under dark conditions (Fig. 6A). However, under continuous white light conditions, the *OsbZIP1.1* expression level was higher than *OsbZIP1.*2 in *OsbZIP1*^OE^ seedlings (Fig. 6A). This points towards the functional significance of the alternate splicing of *OsbZIP1* where the two transcripts have inverse levels under light/dark conditions and may coordinate to regulate seedling development.

To investigate the role of *OsbZIP1* in light-regulated development in rice, we analyzed the *OsbZIP1* transgenic plants under 100 µmol/m^2^/s of continuous white light and dark. When grown under white light, the 5-day-old *OsbZIP1*^OE^ seedlings exhibit a decrease in total height and second leaf length (Fig. 6B-D). In contrast, *OsbZIP1*^KD^ seedlings showed an increase in total seedling height and second leaf length as compared to its vector control (Fig. S10A, B). Marginal changes were observed in the first leaf length of both *OsbZIP1*^OE^ and *OsbZIP1*^KD^ seedlings under both light and dark conditions. No significant difference in the coleoptile length and root length of the seedlings was observed under white light as well as dark conditions (Fig. S11). However, when grown under continuous dark conditions for 5 days, the *OsbZIP1*^OE^ seedlings showed a reduction of 17.64% and 64.5% in the average total seedling height and second leaf length, respectively (Fig.6C, D).

To check if OsbZIP1 differs from AtHY5 in its response to different wavelengths of monochromatic light, the *OsbZIP1*^OE^ as well as *OsbZIP1*^KD^ seedlings were grown under red, blue and far-red light under two different intensities (15 µmol/m^2^/s and 30 µmol/m^2^/s) for 5 days (Fig. 6E). No significant difference was observed in the average coleoptile length and the average first leaf length of the *OsbZIP1*^OE^ seedlings as compared to the vector control under red, blue as well as under far-red light conditions (Fig. S12). However, under 15 µmol/m^2^/s red light, a reduction of 46.7% and 60.5% was observed in the total seedling height and second leaf length, respectively, of the 5-day-old *OsbZIP1*^OE^ seedlings (Fig. 6F, G). A reduction of 61% was also observed in the root length of the *OsbZIP1*^OE^ seedlings (Fig. S13E). A similar trend was seen when *OsbZIP1*^OE^ seedlings were grown under 30 µmol/m^2^/s of red light (Fig. 6E). We observed a decrease of 40.7%, 57.6% and 54.7% in the average total seedling height, second leaf length and total root length, respectively, in the 5-day-old *OsbZIP1*^OE^ seedlings (Fig. 6F, G, S13E).

Under 15 µmol/m^2^/s and 30 µmol/m^2^/s blue light conditions, a reduction was observed in total shoot length, root length and second leaf length of 5-day-old *OsbZIP1*^OE^ seedlings as compared to the vector control (Fig. 6E, F, G, S13A). A decrease in total seedling height and second leaf length of the *OsbZIP1*^OE^ seedlings was also observed under far-red light conditions at both light intensities (Fig. 6E, F, G). Under 30 µmol/m^2^/s blue light conditions, the transcript levels of *OsbZIP1.2* were much higher than *OsbZIP1.1* while *OsbZIP1.2* levels were less than *OsbZIP1.1* at 15 µmol/m^2^/s blue light (Fig. S13C, D). This indicates that *OsbZIP1.1* and its alternative splice form might function at different light intensities; however, it requires more detailed investigation to provide credibility to this observation.

In contrast, under all the three monochromatic light conditions, the total seedling height was higher in the 5-day-old *OsbZIP1*^KD^ seedlings at both the light intensities (Fig. 6E, H). An average increase of 36.3%, 38.11% and 8.26% was observed in seedling height under 15 µmol/m^2^/s red, blue and far-red light, respectively, while an increase of 29.6, 38.41 and 30.83% in seedling height was observed under 30 µmol/m^2^/s red, blue and far-red light, respectively, in the *OsbZIP1*^KD^ seedlings as compared to the vector control. The coleoptile length was, however, unchanged in the *OsbZIP1*^KD^ seedlings as compared to the vector control in these conditions (Fig. S12). The second leaf length was significantly higher in *OsbZIP1*^KD^ seedlings than the vector control; 80.5 % increase under blue light, 96% under red light and 45.5 % under 15 µmol/m^2^/s far-red light conditions (Fig. 6I). A similar trend in increase in second leaf length (44.33% under red light, 72.6% under blue light and 24.7% under far-red light) was observed at 30 µmol/m^2^/s (Fig. 6I). An increase in total root length was also observed in *OsbZIP1*^KD^ seedlings under these monochromatic light conditions (Fig. S13B, F).

Thus, *OsbZIP1* overexpression as well as RNAi transgenic rice plants show drastic change in the total height of the 5-day-old seedlings as well as changes in root length, first and second leaf length in 5-day-old seedlings under blue, red and far-red monochromatic lights.

### *OsbZIP1* expression in rice impacts total height, flag-leaf and Y-leaf length, and flowering time

The *OsbZIP1*^OE^ plants showed a semi-dwarf phenotype at the stage of panicle emergence with about 11% reduction in height compared to the vector control plants (Fig. 7A, 7D). The reduction in total height can be due to reduction in culm length or a reduction in flag leaf length. We observed a significant difference (mean 22.8% reduction) in the culm length of *OsbZIP1*^OE^ plants and vector control (Fig. 7C, S14). Further, the difference in culm length can be due to difference in either the length of the internodes or due to difference in the number of nodes. There was no significant difference in the number of nodes, length of the first internode, second internode and third internode (Fig. S14). However, a significant difference was observed in the length of the second last internode (mean 27.9% reduction) as well as the last internode which bears the panicle (mean 17.6% reduction) (Fig. 7C). Thus, the maximum reduction was observed in the length of the second last internode followed by the last internode.

**Figure 7.**
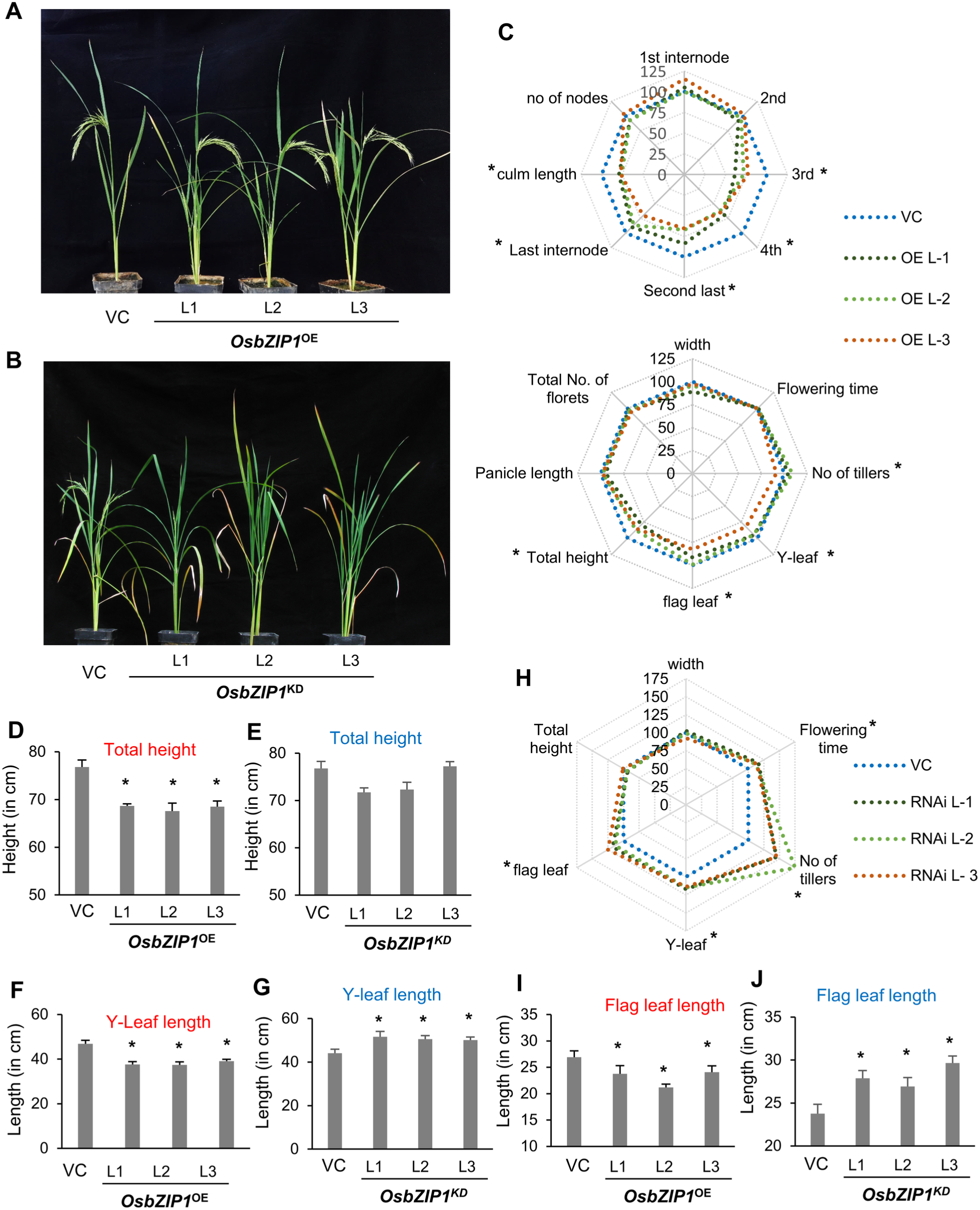
*OsbZIP1* ectopic expression impacts plant height through changes in internode, flag leaf and Y-leaf length. Representative picture shows the phenotype of **(A)** *OsbZIP1*^OE^ and **(B)** *OsbZIP1*^KD^ rice plants at reproductive stage grown in a containment net-house. **(C)** Morphometric parameters of the three *OsbZIP1*^OE^ transgenic lines compared with pb4NU vector control (VC). A percentage of the mean values (n=10) with that of VC set at 100% as reference represents a data point in this plot. (D-E) Histograms show total height (in cm) of the three **(D)** *OsbZIP1*^OE^ and **(E)** *OsbZIP1*^KD^ transgenic lines. Y-leaf length (in cm) of the three **(F)** *OsbZIP1*^OE^ and **(G)** *OsbZIP1*^KD^ transgenic lines shown in histograms. **(H)** Morphometric parameters of the three *OsbZIP1*^KD^ transgenic lines compared with pANDA vector control (VC). Flag leaf length (in cm) of the three **(I)** *OsbZIP1*^OE^ and **(J)** *OsbZIP1*^KD^ transgenic lines shows in histograms. Significance of value by T-test p-value < 0.01 in overexpression/RNAi plants vs respective vector control shown by asterisk (*).

We did not observe significant difference in the average total height of the *OsbZIP1*^KD^ plants and their respective vector control plants (Fig. 7B, E). However, a significant increase in the length of the flag leaf (mean 18.5%) and Y-leaf (mean 15.2 %) was observed in the *OsbZIP1*^KD^ plants as compared to the vector control (Fig. 7G, J). This contrasts with the decrease in Y-leaf and flag leaf length observed in case of the overexpression *OsbZIP1*^OE^ plants in rice (Fig. 7F, I). The average flowering time of the *OsbZIP1*^OE^ plants was found to be similar to the vector control with no significant difference (Fig. 7C). In contrast to the overexpression of *OsbZIP48* in rice that showed reduction in culm length while compromising on the fertility (Burman et al., 2018), no significant difference was observed in the panicle length and average total number of florets of the *OsbZIP1*^OE^ plants and vector control (Fig. 7C). However, the number of days taken for panicle emergence was found to be more in the RNAi *OsbZIP1*^KD^ plants compared to the wild type and vector control. The three *OsbZIP1*^KD^ lines showed an average number of 119.3, 115.6 and 116.9 days to flower, respectively, as compared to the vector control which took an average of 104 days to flower (Fig. 7B, 7H and S10). The panicle length and total number of florets of the *OsbZIP1*^KD^ plants are similar to the vector control (Fig. S10).

Thus, the detailed morphometric analysis of the overexpression and RNAi transgenics revealed that *OsbZIP1* might play a redundant role in the regulation of plant height in rice by affecting the internode length of the second-last and last internode (which bears the panicle) in the rice culm. It may also play a role in the regulation of flag leaf and Y- leaf length as well as flowering time. However, unlike the overexpression of *OsbZIP48*, *OsbZIP1* overexpression does not adversely affect the fertility of the plant while leading to a reduced plant height.

## Discussion

### OsbZIP1 is a functional homolog of AtHY5

Light as an environmental factor alters abundance of transcripts and proteins. Alternative splicing (AS) and ubiquitin dependent protein degradation are two major processes that regulate abundance of key regulators at post-transcriptional and post-translational level, respectively. Increasingly data based upon next generation sequencing have indicated that about 40% - 60% of intron containing genes undergo AS in rice and *Arabidopsis* (Marquez et al., 2012, Dong et. al., 2018). In the present study, we observed that *OsbZIP1* encoding a bZIP domain containing transcription factor undergoes AS such that the two protein isoforms differ in presence of COP1 binding domain while retaining the conserved bZIP domain. OsbZIP1.1 produced from full length (612 bp) transcript, is light-regulated, shows interaction with OsCOP1 and is phosphorylated by OsCK2ɑ3 that regulates its interaction with OsCOP1. It is also present in much higher relative abundance than OsbZIP1.2 which is a product of short (345 bp) alternatively spliced transcript. *OsbZIP1.2* transcript as well as protein levels were quite low and could not be detected on same immunoblots as OsbZIP1.1, thus requiring much higher total protein amount be loaded for its detection (Fig. 1). The relative proportion of *OsbZIP1.2* / *OsbZIP1.1* transcript also do not appear to be drastically different among various samples across light/dark conditions and developmental stages except in root tissues. In comparison to OsbZIP1.1, the levels of OsbZIP1.2 appear to be more stable under light and dark conditions, both at the protein and transcript level (Fig. 2D, 4J, K and 6A). OsbZIP1.2, which differs in its subcellular localization from OsbZIP1.1, also does not show interaction with OsCOP1 and OsCK2ɑ3 (Fig. 3A, 4H, 5B). However, *OsbZIP1.2* is able to complement the *Arabidopsis hy5* mutant in restoring hypocotyl length, chlorophyll content, cotyledon angle, lateral root number and lateral root angle in a way similar to the complementation of the *hy5* by another rice homolog, *OsbZIP48* (Burman et al., 2018) (Fig. 4, S3, S4 and S5). Moreover, both OsbZIP1.1 and OsbZIP1.2 lack an activation domain like OsbZIP48 (Fig. 2E). AtHY5 also does not show any transcription activation in yeast (Ang et al., 1998).

*AtHYH*, a homolog of *AtHY5* in *Arabidopsis,* has four splice variants generated by intron retention and Alt-type AS events (Li et al., 2017). HYH.1, HYH.3 and HYH.4 lack COP1 interaction domain while retaining the bZIP DNA binding domain. *OsbZIP1* also seems to undergo Alt-DA type (alternative donor/acceptor) alternative splicing similar to *AtHYH.1*. All the splice forms of *AtHYH* are also able to complement the elongated hypocotyl phenotype of *Arabidopsis hy5*. Overexpression of *AtHYH* spliced forms in wild type *Arabidopsis* does not affect the hypocotyl elongation, a phenotype similar to overexpression of *AtHY5* and *OsbZIP48* in wild type *Arabidopsis* (Li et al., 2017). While *AtHY5* and *AtHYH* are majorly equivalent in their function, HYH is involved in only blue light mediated development compared to HY5 that functions under all monochromatic light conditions (Holm et al., 2002; Sibout et al., 2006). HY5 also regulates HYH at the protein level, but unlike HY5, HYH is not involved in hypocotyl elongation. Thus, OsbZIP1 which appears to function under all monochromatic light conditions functionally resembles AtHY5 rather than AtHYH. In another similarity with *HY5,* overexpression of *OsbZIP1* in *Arabidopsis* did not show any change in hypocotyl length, cotyledon angle or chlorophyll content. However, overexpression of *OsbZIP1.2* did lead to a photomorphogenic response in *Arabidopsis* under dark conditions. Such over-expression of *AtHY5* lacking the COP1 binding domain in *Arabidopsis* also leads to a hyper-photomorphogenic phenotype under white light as well as monochromatic lights but not in dark (Ang et. al. 1998). Thus, OsbZIP1 can be claimed to be a functional homolog of *AtHY5* but has likely acquired some newer functions.

### Independent yet overlapping functions *of* OsbZIP1.1 and OsbZIP1.2 in photo- and skoto- morphogenesis

Despite its inability of transcriptional activation in yeast, AtHY5 has been reported to act as both activator and repressor of a large number of genes using ChIP-chip analysis (Zhang et al., 2011). This could be due to its interaction with other transcription factors. Since both OsbZIP1.1 and OsbZIP1.2 could independently form homo- and hetero- dimers with OsbZIP48, it is likely that OsbZIP1 also resembles AtHY5 for activation or repression of gene expression in rice (Fig. 3B, C). Further, OsbZIP1.1 localizes in the nucleus and OsbZIP1.2 localizes all over the cell including the nucleus (Fig. 3A). Both these observations indicate that these isoforms may regulate some processes in an overlapping manner and also independently regulate some other processes in the cell. This is also supported by the fact that only OsbZIP1.1 levels are light-regulated while OsbZIP1.2 levels are not much affected by light/dark conditions (Fig. 2D, 4J, K).

Photomorphogenesis in monocots like rice is different from that of dicots like *Arabidopsis* as monocots exhibit partial photomorphogenic response in dark (Zhang et al 2006; Burman et al., 2018). In rice under dark conditions, the first leaf emerges from the coleoptile around 4-5 days after germination while second leaf emerges around 6-7 days after germination. Interestingly, over a time course of 2-10 days in dark, transcript levels of *OsbZIP1.2* with particular peaks at 4^th^, 7^th^ and 10^th^ day seem to coincide with leaf emergence from the coleoptiles of etiolated seedlings (Fig. 2A, B). OsbZIP1.2 might be involved in partial photomorphogenic development under dark as is evident from reduced hypocotyl length phenotype of WT/35S:*OsbZIP1.2* lines in *Arabidopsis* under dark and increased OsbZIP1.2 transcript levels, reduced total seedling height and second leaf length of WT/Ubi: *OsbZIP1* gain-of-function transgenic lines in rice under dark (Fig. 4D, F, 6A, C and D). *OsbZIP1.2* levels might be associated with leaf development under dark conditions as its peaks coincide with days of leaf emergence although this observation needs further validation (Fig.2C). In contrast, *OsbZIP1.1* appears to be associated with early stages of seedling development under light conditions (Fig. 2B). Notably, *AtHY5* levels also peak at around 2-3 days after germination (Hardtke et al., 2000). This is unlike the other homolog OsbZIP48 that shows a similar expression profile under both light and dark conditions and is rather regulated developmentally (Burman et al. 2018).

### Light mediated regulation of OsbZIP1.1 by OsCOP1 and post-translation modification

In *Arabidopsis* under dark conditions, COP1 localizes in the nucleus and binds to HY5, thus causing proteasome-mediated degradation of HY5 in the nucleus (Oyama et al., 1997; Ang et al., 1998; Osterlund et al., 2000), whereas in light, COP1 localizes to the cytoplasm preventing degradation of HY5 (von Arnim and Deng, 1994). The 36 amino acid stretch between residues 25 and 60 in AtHY5 acts as the COP1 binding domain and is involved in the interaction with the C-terminal WD40 domain of AtCOP1. The full-length forms of HY5 homologs in rice, i.e., OsbZIP1, OsbZIP48 and OsbZIP18, possess a conserved COP1 binding domain indicating that COP1 might play a role in their regulation as well. However, OsbZIP48 does not interact with OsCOP1, essentially because of invariant COP1 interaction motif within the domain (Fig. 4I, S6 A). Interaction between OsbZIP1 with OsCOP1 shows that the two HY5 homologs from rice (OsbZIP48 and OsbZIP1) differ from one another in their regulatory mechanisms. Further, analysis of protein levels of OsbZIP1.1 *in vitro* as well as *in vivo* upon light/dark treatments or upon transition of seedlings from light to dark and vice-versa, clearly reflect that OsbZIP1.1 is light regulated. Unchanged levels of OsbZIP1.1 in the *in-vitro* degradation assay when MG132, an inhibitor of 26S proteasome was used, indicate that degradation of OsbZIP1.1 is ubiquitin-26S proteasome dependent (Fig. 4 H, I, J, K). Thus, both OsbZIP1.1 and AtHY5 are dark labile at the protein level and we suggest that OsbZIP1 is more similar to AtHY5 in its regulation by COP1 as compared to OsbZIP48.

The Immunoblot analysis of OsbZIP1.1 revealed the presence of two bands running at an apparent molecular weight around 33-35 kDa with the upper band corresponding to the phosphorylated protein as detected by anti-phosphoserine antibody after *in vivo* immunoprecipitation (Fig. 5D). This indicated the existence of a phosphorylated form of OsbZIP1.1 in a way similar to AtHY5 and OsbZIP48 (Osterlund et al., 2000; Burman et al., 2018). AtHY5 was proposed to be phosphorylated by AtCK2, although no experimental evidence of physical interaction between AtHY5 and AtCK2 was reported (Hardtke et al., 2001). CK2 (CASEIN KINASE 2), a serine/threonine protein kinase exists as a heterotetramer consisting of two catalytic alpha subunits (CK2ɑ) and two regulatory beta subunits (CK2ß) (Hardtke et al., 2001). The regulatory and catalytic subunits can exist independently as monomers both in the nucleus and cytoplasm and exhibit independent functions (Filhol et al., 2004). Rice CK2 was shown to exist as a tetramer of ɑ2/ɑ3 catalytic subunits and ß1/ß3 regulatory subunits. Earlier reports showed that CK2ɑ3/CK2ß3 holoenzyme is able to phosphorylate PT8 (rice phosphate transporter) under Pi (inorganic phosphorous) sufficient conditions, thus playing a role in phosphate homeostasis in rice (Chen et al., 2015).

Recent studies have suggested that AtHY5 is phosphorylated by SPA1 kinase under both light and dark conditions (Wang et al., 2021). It is possible that SPAs might also be involved in the phosphorylation of OsbZIP1.1 although it needs to be experimentally validated. In contrast phosphorylation of OsbZIP1.1 by CK2ɑ3 appears to be only under light conditions (Fig 2D, 5D, 4K). OsbZIP48, although exists as a phosphorylated form *in vivo*, does not get phosphorylated by OsCK2ɑ3 (Fig. S7). Thus, OsbZIP1.1 is post-translationally regulated in a manner more similar to AtHY5 as compared to OsbZIP48.

The significance of phosphorylation in case of OsbZIP48 needs to be understood further while in case of OsbZIP1.1 phosphorylation possibly acts as a mechanism of protection from degradation. While OsCOP1 shows a differential binding affinity towards the phosphorylated and unphosphorylated forms of OsbZIP1, interacting more strongly with the unphosphorylated form as compared to the phosphorylated form; it does not interfere with the phosphorylation of OsbZIP1 (Fig. 5). The phosphorylation alters affinity of OsbZIP1 towards OsCOP1 preventing it from being degraded under the light conditions (Fig. 5). The unphosphorylated OsbZIP1.1 having higher binding affinity to COP1 gets degraded much faster under dark conditions where OsCOP1 accumulates in the nucleus. In light, OsCOP1 might move out of the nucleus and it can be speculated that the unphosphorylated OsbZIP1.1 might interact more strongly with its target promoters as in case of AtHY5, although it remains to be experimentally validated. This small pool of phosphorylated OsbZIP1.1, which is less susceptible to degradation by OsCOP1, might also be physiologically less active similar to AtHY5 (Wang et al 2021; Hardtke et al 2000). Notably, upon transition of rice seedlings from dark to light, while increase in OsbZIP1.1 level is observed within 5 hours, it takes around 10-15 hours for the phosphorylated OsbZIP1.1 (upper band) to appear on the immunoblot (Fig. 4K). This indicates an additional set of regulation through OsCK2ɑ3 that is triggered once OsbZIP1.1 is present in an abundant quantity under continuous light conditions. Thus, the regulation of OsbZIP1.1 activity takes place by two processes -- one, by its physical interaction with OsCOP1, which might depend on the light mediated nuclear exclusion of OsCOP1 and second, through phosphorylation by serine/threonine protein kinase, OsCK2, whose activity is known to be light dependent and which modulates OsbZIP1’s ability to interact with OsCOP1. Thus, both the mechanisms of regulation of OsbZIP1.1 activity influence its interaction with OsCOP1. In contrast, OsbZIP1.2 that lacks a COP1 binding domain, does not physically interact with either OsCOP1 or OsCK2 and exists as a single unphosphorylated form in the immunoblots, and possibly imparts it biological activity both in dark and light. The levels of OsbZIP1.2 stay largely unaltered between dark and light transitions (Fig. 4K). Further, it can interact with OsbZIP48 or form homodimers independently and also promote photomorphogenic response under dark when *OsbZIP1.2* is overexpressed in *Arabidopsis*. Thus, it is possible that OsbZIP1.2 with intact bZIP domain may also regulate function of downstream target genes involved in light signaling singularly or synergistically with OsbZIP1.1. This in all probability indicates that while OsbZIP1.1 is key regulator of photomorphogenesis in light whereas OsbZIP1.2 regulates the partial photomorphogenesis in rice under dark conditions.

### Neofunctionalization of *OsbZIP1* and its impact on light responses in rice

HY5 is known to function downstream to multiple photoreceptors (such as phyA, phyB, cryptochromes and UV-B) acting as a positive regulator of photomorphogenesis under all the wavelengths of light (Koornneef et al., 1980; Ang and Deng, 1984; Oyama et al., 1997; Ulm et al., 2004). Recently, it has been shown that all the three AtHY5 homologs in rice, *OsbZIP1*, *OsbZIP18* and *OsbZIP48* are induced by UV-B irradiation with *OsbZIP1* showing ̴6 folds upregulation (Sun et al., 2022). We observed that overexpression of *OsbZIP1* in rice leads to a reduction in seedling height and knockdown expression lead to elongated seedlings when grown under continuous white, red, far-red and blue light conditions (Fig. 6). This phenotype is primarily manifested through changes in second leaf length under all these conditions, whereas coleoptile length and first leaf length remained largely unaffected by changes in *OsbZIP1* expression. The phenotypic responses of *OsbZIP1^OE^* and *OsbZIP1^KD^* lines clearly suggest that OsbZIP1 acts downstream to multiple photoreceptors including phytochromes and cryptochromes. Further, the differential expression of both transcripts of *OsbZIP1* under different light intensities and its downstream effect needs further investigation. At present, it is tempting to speculate that *OsbZIP1.1* may work under low light intensities while *OsbZIP1.2* may function under high light intensity, thereby helping the plant adapt to different intensities of light. Similar observations have been reported for *CCA1* and *SPA3*. The circadian clock regulator *CCA1* has an alternatively spliced variant CCA1ß which accumulates under high light conditions (Shang et al., 2017). The altD type of alternative splicing of *SPA3* is also promoted by light (Cheng et al., 2018).

Previously, we showed that OsbZIP48 binds directly to the promoter of *OsKO2*, which encodes a gibberellin biosynthesis pathway enzyme *ent*-kaurene oxidase 2. Since both OsbZIP1.1 and OsbZIP1.2 homodimerize as well as heterodimerize with OsbZIP48, it can be speculated that they form a complex with OsbZIP48 that binds to promoter of these gibberellin biosynthesis genes and thereby altering their expression leading to a semi-dwarf phenotype. Accordingly, we observed that the *OsbZIP1* levels were upregulated almost 9 folds in the OsbZIP48^OE^ plants (Burman et al., 2018). It is also possible that they might function redundantly since the percentage decrease in culm length observed in case of *OsbZIP1^OE^* plants is less than that of *OsbZIP48^OE^*. Also, overexpression of OsbZIP1 in rice does not impact panicle length, total number of fertile florets or percentage fertility in contrast to *OsbZIP48^OE^* rice plants that showed a significant decrease in above mentioned agronomic traits (Burman et al., 2018). Further, *OsbZIP1*^KD^ plants do not show significant change in the average total height of the three transgenic lines as compared to the vector control (Fig. 7). Recently, characterization of *OsbZIP1/our1* mutants in rice did not reveal any change in height of mature plants when compared to the wild type (Hasegawa et al., 2021). Also, the OsbZIP18 overexpression in rice has been reported to cause a significant reduction in plant height, tiller number and severe dark brown pigmentation in mature leaves (Sun et al., 2022). Thus, all the three AtHY5 homologs in rice regulate plant height and may exhibit redundancy. However, it requires further investigations if *OsbZIP48* and OsbZIP18 can independently regulate plant height when *OsbZIP1* is downregulated.

OsbZIP1 functions in seedling development as well as in mature plant development in response to light and has acquired some novel functions in comparison to both OsbZIP48 and AtHY5. All these homologs of AtHY5 in rice may have overlapping yet distinct functions. A recent study from our laboratory showed that *OsbZIP48* and *OsbZIP18* are differentially expressed under high temperature conditions in rice cultivars with contrasting heat stress tolerance, whereas *OsbZIP1* levels were unaltered under above mentioned conditions (Sharma et al., 2021). *OsbZIP18* was also found to be induced by nitrogen deficiency and polymorphisms in its promoter affects branched chain amino acids levels in rice (Sun et al., 2020). A recent study suggested that *OsbZIP1* suppresses auxin signaling by altering the expression of auxin-responsive genes to control root architecture (Hasegawa et al. 2021). Thus, HY5 homologs in rice have undergone neofunctionalization with increased repertoire of effects in response to environmental and developmental clues. Being transcription factors, how this functional diversification among HY5 homologs -- *OsbZIP48, OsbZIP1 and OsbZIP18 --* is manifested in regulating expression of downstream target genes remains to be understood in greater detail.

## Materials and methods

### Plant materials and quantitative RT-PCR analysis

Rice seeds (Oryza sativa var. *indica*) procured from Indian Agricultural Research Institute (IARI), New Delhi were subjected to surface sterilisation by treating with 70% ethanol for 30s followed by 0.1% HgCl_2_ for 10 min and washed three times with autoclaved Milli-Q water. These seeds were then incubated for 16 h at 28±1°C and were grown hydroponically for 10 days on Yoshida medium in a culture room maintained at 28±1°C under white light of 100 µmol/m^2^/s provided by Cool Daylight fluorescent lamps (Philips; TL 5800°K) (Burman et al. 2018). Seedlings were harvested every 24 hrs starting from day 2 until day 10 and frozen in liquid nitrogen followed by storage at -70°C. Tissue for developmental stages was harvested from rice plants grown in the fields of IARI as per the following description for stages: SAM: up to 0.5 mm, panicle stages: P1-1: 0.5-2 mm, P1-2: 2-5 mm, P1-3: 5-10 mm, P1: 0-3 cm, P2: 3-5 cm, P3: 5-10 cm, P4: 10-15 cm, P5: 15-22 cm, P6: 22-30 cm, seed stages: S1, 0–2 DAP; S2, 3–4 DAP; S3, 5–10 DAP; S4, 11–20 DAP; and S5, 21–29 DAP (Jain et al., 2007, Sharma et al., 2012). For RT-qPCR analysis of *OsbZIP1*^OE^ and *OsbZIP1^KD^*, seedlings were grown on half strength Murashige and Skoog (MS) medium with 1% Phytagel in continuous 100 µmol/m^2^/s of white light, 15 and 30 µmol/m^2^/s of blue light and also in dark, as described above, for 5 days (Zhang et al. 2006). Total RNA was extracted from the harvested tissue using TRIzol reagent (Invitrogen, USA) as per the manufacturer’s instructions. Transcriptor First-Strand cDNA Synthesis Kit (Roche, USA) was used for synthesis of cDNA using random hexamer primers. RT-qPCR was performed using LightCycler 480II Real Time system (Roche, USA) as per the manufacturer’s instructions. *UBIQUITIN5* was used as an internal standard relative to which the mRNA levels of *OsbZIP1.1* and *OsbZIP1.2* were computed. Three technical and three biological replicates were used for each sample. The conditions used for RT-qPCR have been described previously (Burman et al., 2018). 2^−ΔCt^ method was used to calculate the relative expression with respect to *UBIQUITIN 5*and relative fold change was calculated with respect to vector control (Livak et al., 2001). List of primers is given in Table S1.

### Rapid amplification of cDNA ends (RACE) and site-directed mutagenesis

The cDNA prepared from 10-day-old *OsbZIP48^OE^* seedlings was used to perform 5’ and 3’ RACE using SMARTer® RACE 5’/3’ Kit (Clontech, TaKaRa Bio, USA) as per manufacturer’s instructions followed by Sanger based sequencing of the RACE products. Site-directed mutagenesis was performed using Phusion DNA polymerase PCR mix (New England Biolabs, UK). Primers were designed specifically to replace the nucleotides coding for serine at the 44^th^ position in the CDS of OsbZIP1.1 with those for alanine as given in table S1.

### Immunoblotting

For immunoblot analysis, protein was extracted from tissue harvested from different stages of seed and panicle development as described above. Light to dark and dark to light transition experiments were performed as described by Burman et. al. (2018). For checking the levels of OsbZIP1.1 and OsbZIP1.2 in light and dark, the *OsbZIP1*^OE^ rice seeds were surface-sterilized and grown on NB medium (Himedia, India, cat no. PT107) at 32°C for 7 days in 100 µmol/m^2^/s of white light or kept in dark room at 28°C for initiation of calli. The tissue was homogenized in a buffer {200 mM Tris, pH 8, 100 mM NaCl, 10 mM EDTA, 10 mM DTT, 5% glycerol, 0.05% Tween 20, and protease inhibitor cocktail (Sigma, USA) and centrifuged at 4°C. The supernatant was used to estimate the protein concentration using Bradford assay (Bradford, 1976). Immunoblotting was performed by loading equal concentrations of protein extracts on SDS-PAGE gel (Sharma et al., 2014). A 1:3000 dilution of custom-made peptide specific antibody was used for detection of OsbZIP1.1 and OsbZIP1.2 and actin was used as a loading control detected by anti-actin antibody (ThermoFisher Scientific, USA, catalog no. MA1-744).

### Subcellular localization and BiFC

For BiFC and subcellular localization, particle bombardment was performed using the Biolistic PDS-1000/He particle delivery system (Bio-Rad, USA) according to the protocol described earlier (Sharma et al., 2022). After particle bombardment of the onion peel cells, they were kept at 28±1°C in dark for 16 h and observed under confocal microscope (Leica microsystems, TCS, SP5). BiFC for each combination was repeated twice and appropriate negative controls were used (Kudla and Bock, 2016).

### Yeast transactivation and two-hybrid assay

OsbZIP1.1 and OsbZIP1.2 CDS were cloned in pDEST-GBKTT7 (CD3-764) obtained from TAIR (https://www.arabidopsis. org) and transformed in the AH109 yeast strain (Clontech, TaKaRa Bio, USA). Yeast transformation was performed as described in the Yeastmaker Yeast Transformation System 2 User Manual (Clontech, TaKaRa Bio, USA). Serial dilution of transformed colonies were dropped on –Trp/-His synthetic dropout selection medium. For yeast two-hybrid assay, full length CDS of OsbZIP1.1, OsbZIP1.2, OsCK2ɑ2, OsCK2ɑ3, OsCK2ß1 and OsCK2ß2 were cloned in pDEST-GBKTT7 and pDEST-GADT7 vectors as required and transformed in yeast AH109 strain as described above. The serially diluted constructs were dropped on -His/-Leucine/-Trp dropout selection media. The sealed plates were incubated at 30°C for 3 to 6 d.

### *In vitro* pull-down assay

6X-His tagged OsbZIP1.1 and OsbZIP1.2 as well as GST tagged OsCOP1 and OsCK2 were expressed in *E. coli* [C41 pLysS (Lucigen, USA)] by cloning their coding sequences in pET28a(+) vector (Novagen, Netherlands) and pGEX4T1 vector (Amersham Biosciences, USA), respectively. *In vitro* pull-down assay was performed as described earlier (Maitra- Majee et al., 2020) by immobilizing GST, GST-OsCOP1 and GST-OsCK2ɑ3 onto GSH-Sepharose (GE Healthcare, USA) by mixing at 4°C for 30 min followed by centrifugation and washing with PBS buffer (Kepinski, 2009). Protein extracts of 6X-His tagged OsbZIP1.1 and OsbZIP1.2 were incubated with the immobilised protein for 1h at 4°C in elution buffer (150 mM NaCl, 100 mM Tris–Cl-pH 7.5, 0.5% NP40, protease inhibitor, MG132). Proteins were eluted from the beads and subjected to western blotting as described earlier (Sharma et al., 2014).

### *In vitro* degradation assay

*In vitro* degradation assay was performed as described by Thakur et al. (2005). Total protein was isolated from 4-day-old light- and dark-grown seedlings by grinding them in a buffer (25 mM Tris, pH 7.5, 10 mM MgCl_2_, 5 mM DTT, 10 mM NaCl and 15 mM ATP). 1 µg 6X-His- OsbZIP1.1 protein induced in bacteria was incubated with equal amounts of the light- and dark-grown seedling protein extract at 30°C for the indicated time points. The reactions were stopped by adding 5X loading dye and samples resolved on SDS-PAGE followed by western blotting. The membrane was probed with 1:10000 anti-His antibody (Sigma, USA) followed by 1:20000 anti-mouse antibody (Sigma, USA) and the signal was detected using Luminata Forte Western reagent (Milipore, USA).

### *In vitro* kinase assay and *In vivo* immunoprecipitation

The His-tagged proteins were purified using Ni-NTA and GST-tagged proteins were purified using GST- Sepharose as per manufacturer’s instructions. For *in-vitro* kinase assay, 1 µg 6X- His-OsbZIP1.1 protein induced in bacteria and purified was mixed with 500 ng purified GST- OsCK2ɑ3 and kinase buffer (20 mM Tris, 1 mM MgCl₂, 0.5 mM CaCl₂ and 2 mM DTT) in the presence of either 0.5 mM ATP or 0.5 mM GTP and incubated at 30°C for 30 min. The GST protein induced in bacteria was used as negative control. 10 µM DRB and 400 U of lambda phosphatase were used as inhibitor and for dephosphorylation, respectively. The reactions were stopped by adding 5X loading dye and protein samples subjected to SDS-PAGE followed by western blotting. The membrane was probed with 1:10000 anti-phosphoserine or anti-threonine antibody followed by 1:20000 anti-mouse antibody and detected using Luminata Forte Western reagent (Millipore, USA). Same protocol was followed for in vitro kinase assay using the mutated osbzip1.1 protein. For *in vivo* immunoprecipitation, 5-day-old light- and dark-grown seedling protein extract was incubated with 1 µl anti-phosphoserine antibody (Sigma, USA) with gentle rotation for 2h at 4°C followed by centrifugation and subjecting the pellet to western blotting as described by Matsushita et al. (2013).

### Generation of rice and *Arabidopsis* transgenics and their morphometric analysis

Floral dip method was used to raise *Arabidopsis* transgenics (Clough and Bent, 1998). *hy5* mutant seeds (SALK_096651C) were obtained from the Arabidopsis Biological Resource Center (https://abrc.osu.edu/). The *Arabidopsis hy5* mutant was transformed with the 780 bp genomic region of *OsbZIP1* cloned in a binary overexpression vector through *Agrobacterium*-mediated transformation. Five transgenic lines of each construct were checked for the presence of the transgene by PCR for the marker hygromycin resistance gene (Fig. S2). Morphometric analysis of *Arabidopsis* transgenics including hypocotyl length measurement, chlorophyll estimation, cotyledon angle measurement, lateral roots analysis were performed as described in Burman et al. (2018) and Maitra-Majee et al. (2020).

Rice transgenics were generated using tissue culture protocol as described by Toki et al. (2006) with modifications as described by Burman et al. (2018). For RNAi, 200 bp region from the 3’ CDS of *OsbZIP1* common between both *OsbZIP1.1* and *OsbZIP1.2* was chosen and assessed for siRNA and off-target effects. Morphometric analysis of rice transgenics was done as per IRRI guidelines (http://books.irri.org/getpdf.htm?book=971104000X). For internode measurement, the first internode was designated as the one closest to the root and the panicle bearing internode was considered to be the last. The *OsbZIP1*^OE^ and *OsbZIP1^KD^* seedlings were grown under 15 and 30 µmol/m^2^/s of white light, red, far-red and blue monochromatic light conditions as well as dark for 5 days and morphometrically analysed as described in Zhang et al. (2006).

### Statistical analyses

The data were analyzed by one-way ANOVA followed by T-test, P-value < 0.01 for significant differences.

### Accession Numbers

AtHY5: At5g11260 OsbZIP1:

LOC_Os01g07880, Os01g0174000:

OsbZIP48: LOC_Os06g39960, Os06g0601500;

OsbZIP18: LOC_Os02g10860, Os02g0203000.

## Supplemental Data

**Fig. S1.** Expression analysis of *OsbZIP1.1* and *OsbZIP1.2*.

**Fig. S2**. Verification of transgenic lines by PCR using *HPTII* gene in *Arabidopsis*.

**Fig. S3**. Complementation of *Arabidopsis hy5* mutant by *OsbZIP1* (gDNA) and *OsbZIP1.2* and their overexpression in wild type *Arabidopsis* showing hypocotyl length and chlorophyll content.

**Fig. S4.** Complementation of *Arabidopsis hy5* mutant by *OsbZIP1.2* and *OsbZIP1* (gDNA) and their overexpression in wild type *Arabidopsis* showing chlorophyll content and cotyledon opening angle.

**Fig. S5.** Analysis of Arabidopsis *hy5* mutant and *OsbZIP1* complementation lines for lateral roots number and angle.

**Fig. S6.** Non-interaction of OsbZIP48 with OsCOP1 and interaction of OsbZIP1.1 with CASEIN KINASE 2 subunits a3 (CK2ɑ3) and ß1 (CK2ß1).

**Fig. S7.** Phosphorylation status of OsbZIP48 and its non-interaction with OsCK2α3.

**Fig. S8.** Phosphorylation of OsbZIP1.1 and effect of mutation of its predicted phosphorylation site on its interaction with OsCOP1.

**Fig. S9.** Confirmation of rice overexpression *OsbZIP1*^OE^ and *OsbZIP1*^KD^ RNAi transgenics.

**Fig. S10.** Morphometric analysis of RNAi transgenics of *OsbZIP1* (*OsbZIP1*^KD^) in rice.

**Fig. S11.** Phenotypic analysis of 5-d-old white light and dark grown *OsbZIP1*^OE^ and *OsbZIP1*^KD^ rice seedlings.

**Fig. S12**. Phenotypic analysis of 5-d-old *OsbZIP1*^OE^ and *OsbZIP1*^KD^ rice seedlings grown under red, blue and far-red light.

**Fig. S13.** Root length analysis of *OsbZIP1*^OE^ and *OsbZIP1*^KD^ seedlings under blue and red light and transcript levels of OsbZIP1.1 and OsbZIP1.2 under blue light.

**Fig. S14.** Agro-morphological analysis of overexpression transgenics of *OsbZIP1* (*OsbZIP1*^OE^) in rice at maturity.

**Table S1**. List of primers used in this study and sequence of peptide used for production of anti-OsbZIP1 antibody.

## Acknowledgements

This work was financially supported by the Department of Biotechnology of the Government of India (No. BT/ AGIII/CARI/01/2012) and J.C. Bose National Fellowship grant by the SERB to JPK (SB/S2/JCB-13/2013). The infrastructural support provided by the University Grants Commission, New Delhi (UGC-SAP) and the Department of Science and Technology (FIST and PURSE grants) is gratefully acknowledged. AB and ES acknowledge the fellowship/research associateship awarded by the Council of Scientific and Industrial Research (CSIR), Government of India. NB acknowledges DST for DST-INSPIRE faculty award (DST/INSPIRE/04/2017/001769).

## Notes

### Competing Interest Statement

The authors have declared no competing interest.

